# Robust height-diameter allometries for 41 European tree species: stand characteristics and structure matter

**DOI:** 10.1101/2025.09.29.679156

**Authors:** Natacha Savine, Thomas Cordonnier, Gauthier Ligot, Patrick Vallet

## Abstract

Tree height is a key variable for assessing forest functioning and resources (e.g volume, carbon stocks) at both tree and stand levels. However, direct field measurements are costly and time-consuming. Developing accurate and unbiased height-diameter allometries applicable over large spatial scales is therefore crucial for forest research and management.

Based on the French National Forest Inventory, with 269,460 tree-height observations on 49,120 plots measured, we developed generalized, species-specific height-diameter allometries that integrate stand dendrometrical characteristics and stand structure. The models cover 41 European tree species and the four main stand structures: even-aged, uneven-aged, coppice-with-standards and coppice. To enhance local accuracy, we developed an optional recalibration method at the plot level, assessing how many additional trees should be measured and which ones to select to maximize the improvement of the model. By integrating basal area as stand density and quadratic mean diameter as an indicator of stand development stage, our models enable us to assess the influence stand dendrometrical characteristics on these allometries and evaluate how different species responded to competition.

Results indicate that with increasing competition, tree height tends to be higher for a given diameter, and that stand structure significantly influences 28 out of the 41 species. Local recalibration showed that measuring just one to six trees (among the largest and thinnest diameter) per plot reduced prediction error by 10 to 70%, depending on species.

This study provides a useful, robust and scalable tool for forest research and management, for the most widespread species in Europe, while offering precision for local applications.

## 1. Introduction

Forest management involves not only assessing existing resources but also evaluating how they have changed, or will change, over time. In this respect, along with diameter, tree height is a key variable to assess forest ecosystem functioning and forest ecosystem services such as carbon storage or wood production (Hunter et al., 2013; Teobaldelli et al., 2009). Studying height variability across individuals can be key to understanding important ecological processes, such as competitive and facilitation interactions between individuals.

However, accurately measuring tree height is often time-consuming and technically challenging especially on slopes or stands with low visibility. As a consequence, tree height measurements are often performed on a limited sample of trees.

In biological organisms, allometries defines how the size of an organ changes in relation to the size of another one, or to the size of the whole organism. These allometries are used to understand how variables are interconnected, and can serve as an alternative for variables that are difficult to measure directly. In forestry, allometric models can be used to estimate tree characteristics such as biomass (Pilli et al., 2006) or volume (Vallet et al., 2006), being common examples. For these two allometric models, the most accurate models include height measurements, or height estimates. Tree height can also be estimated indirectly through allometric relationships, most commonly by using diameter at breast height (dbh), a variable that is much easier to obtain in the field (Ciceu et al., 2023).

Height-diameter allometries are known to differ across species, due to biological constraints, limitations and growth strategies inherent to each species (Vázquez-Veloso et al., 2025). Allometries can therefore be species-specific (Adame et al., 2008; Ciceu et al., 2020; Huang et al., 1992) or can take species characteristics directly into account in the model (Sharma et al., 2019b).

It is also essential to consider the specific environmental conditions influencing individual growth. When trees grow in proximity to other individuals, they compete for resources in different ways. In dense areas, where space is limited, trees may allocate more resources to vertical growth, which increases height, rather than to radial growth. This phenomenon can be integrated into allometric models by considering the social status of the tree (Aussenac et al., 2023; Trouvé et al., 2015). Social status can be used per se in the models, or can be included as a competitive index as in Mugasha et al. (2019).

The dendrometrical characteristics of the stand also play a role in shaping height-diameter allometries. Stand density variables such as basal area, for example, are commonly used to account for competition. As a result, trees growing in highly competitive environments present greater height for a given diameter compared to those in less competitive ones.

Ciceu et al (2023) highlighted that dominant height is used to reflect the effect of stand developmental stage in most studies working on height-diameter allometry. However, to obtain this variable, the height of a subset of trees must be measured. The widespread inclusion of dominant height in empirical models is likely due to the strong correlation between dominant height and individual tree heights. Alternative variables, such as the quadratic mean diameter, also reflect stand development stage (Aussenac et al., 2023; Ciceu et al., 2020; Feldpausch et al., 2011; Gonçalves, 2022; Mugasha et al., 2019; Temesgen and v. Gadow, 2004).

The structure of the stand (even-aged, uneven-aged and other configurations) also influences height-diameter allometry(Gonçalves, 2022). Different stand structures reflect different stand dynamics and result in diverse tree diameter distributions. For instance, even-aged stands are characterized by individuals of similar ages and heights, whereas uneven-aged stands exhibit more diverse ages and height structures, potentially resulting in different allometric patterns.

Throughout the diversity of environmental conditions and forest structures, the French metropolitan forest area is covered by 58% of even-aged stands. This leaves 42% of the stands that are managed under other regimes such as uneven-aged, coppice or coppice-with-standards systems (described in figure S1, Supplementary Materials). The even-aged structure has been the focus of most studies on forest structure, as it is one of the most represented and constitutes a simple silvicultural system (Bettinger et al., 2017). However, since structure plays an important role in growth, especially with different competitive relationships, it is necessary to study how height-diameter allometries are affected by each structure specifically. This could prevent overestimations or underestimations when assessing forest resources (Gonçalves, 2022).

Eventually, allometric models can be developed at different scales, from local to continental. While large-scale models can provide higher genericity, they may introduce estimation bias when applied to specific local conditions. Studies by Sullivan et al. (2018), Mensah et al. (2018) and Kearsley et al. (2017) have shown that locally calibrated models perform better than large scale models. To improve the genericity and performance of these models, it can be beneficial to incorporate local factors such as soil fertility, water availability or climate, thus allowing the model to adapt to specific conditions.

In this study, we developed height-diameter allometries for 41 species. We hypothesized that this relationship would be influenced by individual-tree and stand characteristics such as stand basal area, mean diameter and stand structure (even-aged, uneven-aged, etc.). We attempted to provide the most practical and generic models possible, for the most species, and for different stand structures. We also developed a method to locally adapt large-scale models, with as few field measurements as possible.

## 2. Material

### 2.1. Measurement protocol

We used data collected by the French National Forest Inventory (NFI) between 2005 and 2022 (IGN, 2023). The plots were located throughout the French metropolitan area, on a systematic sampling grid of 1×1 km. Each year, one twentieth of the grid was sampled, representing approximately 6000 plots per year. The plots are composed of three concentric subplots with 6-, 9- and 15-meter radii. Very small trees (dbh [7.5-12.5cm [ ) and small trees (dbh [12.5-22.5cm [ ) are measured in the 6-meter radius subplots, medium trees (dbh [22.5 – 37.5cm [ ) are measured in the 9-meter radius subplots, and large (dbh [37.5 – 52.5 cm [ ) and very large trees (dbh over 52.5 cm) are measured in the 15-meter radius subplots. To account for the concentric circle sampling design, the NFI assigns a weight to each tree based on its diameter class (very small, small, medium, large and very large). These weights are then used to calculate stand-level variables, such as quadratic mean diameter and basal area.

Until 2013, height was measured for six trees per diameter class and per species. From 2014 to 2015, only one tree was measured per category and per species. In 2016, the very-small-tree category was added and additional height measurements were carried out on very small trees. During the inventories, height and diameter are measured, along with some other tree variables such as the vegetative state (living or dead) or stem singularities (broken, abnormal, pollarded). The complete measurement protocol is described in Dalmasso et al. (2014).

From 2005 to 2013, the NFI classified stand structures in five categories based on the characteristics of the stand, combining the regeneration method (coppice, coppice-with-standards, even-aged forest, uneven-aged forest, no structure) and vertical distribution. From 2014 to 2022, the NFI used different stand structure classifications, which we aggregated to distinguish four stand structures: even-aged (EA), uneven-aged (UA), coppice-with-standards (CWS) and coppice (C) (aggregation criteria are described in Supplementary Materials, table S1).

### 2.2. Dataset sub-samples

We selected plots dominated by one species and for which stand structure was available. The dominant species had to represent at least 75% of the plot basal area, considering only living trees. We discarded plots for which stand structure was not available (structure not registered, undefined, or registered as no structure). Within the selected plots, we excluded trees with special stem characteristics, such as broken or pollarded trees. This eliminated 1.38% of the dataset. Additionally, we selected species for which there were at least one hundred tree records. Only the combinations of species and stand structure for which we had at least ten tree records were analyzed.

Our sample contained tree records for 51 species in all: 30 species in all four stand structures, 16 species in three structures (14 not present in coppice stands, 2 not present in uneven-aged stands), and five species in two stand structures (even-aged and coppice-with-standards standards). We also discarded the few tree records of coniferous species in coppice stands.

Our final models were fitted for 41 out of the 51 species. For ten species, the fitting procedure did not reach convergence, most likely because we had too few observations or because individual height was not evenly distributed across diameter ranges.

The final calibration dataset contained 269,460 measured trees on 49,120 temporary plots. All 41 species are presented in the Supplementary Materials, table S2, and the ten most sampled species are described in table 1.

**Table 1:** Description of the sampling characteristics of the ten most sampled species in the calibration dataset. EA: Even-aged, UA: Uneven-aged, CWS: Coppice-With-Standards, C: Coppice. The full list is available in Supplementary Materials, Table S1.

| Species | Number of trees | Number of plots | Number of trees per structure |  |  |  | DBH (cm) |  |  | Height (m) |  |  |
| --- | --- | --- | --- | --- | --- | --- | --- | --- | --- | --- | --- | --- |
|  |  |  | EA | UA | CWS | C | min | mean | max | min | mean | max |
| *Abies alba* | 19,055 | 2,361 | 12,502 | 3,772 | 2,781 | 0 | 7.48 | 36.91 | 129.23 | 2.4 | 21.90 | 46.2 |
| *Castanea sativa* | 14,334 | 2,670 | 2,993 | 417 | 7,050 | 3,874 | 7.48 | 27.46 | 262.92 | 2.2 | 15.40 | 34.8 |
| *Fagus sylvatica* | 28,135 | 4,402 | 16,437 | 1,302 | 9,201 | 1,195 | 7.48 | 33.52 | 174.12 | 1.3 | 20.21 | 47.1 |
| *Picea abies* | 19,176 | 2,664 | 14,933 | 2,127 | 2,116 | 0 | 7.51 | 32.14 | 127.01 | 1.6 | 21.68 | 48.4 |
| *Pinus pinaster* | 20,695 | 4,075 | 16,204 | 551 | 3,940 | 0 | 7.48 | 30.61 | 111.09 | 2.6 | 16.93 | 39.1 |
| *Pinus sylvestris* | 21,865 | 3,572 | 15,409 | 1,988 | 4,468 | 0 | 7.48 | 27.22 | 98.96 | 1.7 | 13.97 | 41.2 |
| *Pseudotsuga menziesii* | 13,691 | 2,151 | 11,809 | 207 | 1,675 | 0 | 7.51 | 33.20 | 140.69 | 2.0 | 23.62 | 49.4 |
| *Quercus petraea* | 31,927 | 5,772 | 16,779 | 575 | 13,370 | 1,203 | 7.48 | 35.64 | 137.35 | 1.9 | 20.95 | 44.0 |
| *Quercus pubescens* | 20,689 | 4,655 | 5,527 | 879 | 9,188 | 5,095 | 7.48 | 20.80 | 192.26 | 1.8 | 11.39 | 30.8 |
| *Quercus robur* | 28,049 | 5,691 | 13,362 | 935 | 12,762 | 990 | 7.48 | 35.68 | 164.12 | 2.2 | 18.83 | 43.2 |

### 2.3. Validation datasets

In addition to our calibration dataset, we used two other independent datasets for validation purposes. These datasets enabled us to test both the performance and the robustness of our models, and were not used for the calibration. To assess prediction bias, we conducted a t-test on the residuals (observed minus predicted heights) to determine whether the mean of the prediction error differed significantly from zero.

The internal validation dataset contains NFI data collected in 2023 (IGN, 2023). This data was collected following the same protocol as the calibration dataset. It includes a large number of height measurements from a large number of species and for the various stand structures. By applying the same criteria as for the calibration dataset, we selected 8,771 height measurements from 2,548 plots, covering 40 species. This second dataset will be referred to as the NFI_2023 dataset, and is described in Supplementary Materials, table S3.

We also used an external dataset for validation, which included six species. This data was collected following a different protocol than that of the NFI. The trees were measured between 1920 and 1955. The plots were sampled among pure highly-productivity even-aged permanent plots established by the former French Forest Administration. The measurement and selection protocol are described in Vallet et al. (2006). This third dataset will be referred to as the FFA dataset, and is described in table 2.

**Table 2:** Description of the French Forest Administration validation dataset.

| Species | Number of trees | Number of plots | dbh (cm) |  |  | Height (m) |  |  |
| --- | --- | --- | --- | --- | --- | --- | --- | --- |
|  |  |  | min | mean | max | min | mean | max |
| *Abies alba* | 762 | 28 | 7.64 | 31.99 | 102.81 | 9.9 | 23.45 | 39.4 |
| *Fagus sylvatica* | 1,293 | 50 | 2.86 | 24.54 | 79.58 | 5.4 | 22.16 | 42.5 |
| *Pinus sylvestris* | 389 | 17 | 6.68 | 22.40 | 53.16 | 9.0 | 19.18 | 28.5 |
| *Picea abies* | 309 | 8 | 15.28 | 32.27 | 70.98 | 10.9 | 26.45 | 41.3 |
| *Pseudotsuga menziesii* | 224 | 7 | 4.77 | 14.95 | 34.06 | 5.5 | 15.12 | 25.0 |
| *Quercus petraea* | 1,222 | 58 | 4.46 | 23.15 | 89.45 | 6.0 | 20.55 | 39.5 |

## 3. Method

### 3.1 General principles of the study

Following the general principles for fitting allometric relationships (Mehtätalo et al., 2015; Zhou et al., 2023), we developed generic models: for each species, the same equation was calibrated, incorporating both tree social status (*dbh*/*D*_*g*_) and stand variables. Stand basal area (BA) was used as a proxy for stand density, and quadratic mean diameter (D_g_) for development stage. We also included the effect of stand structure (even-aged, uneven-aged, coppice-with-standards and coppice) on the height-diameter allometry. Our models were tested for accuracy and robustness on both validation datasets.

Finally, to improve the height estimates for a given forest plot, we developed a local recalibration method based on a limited number of height measurements.

### 3.2 Functions

#### 3.2.1 General equation

Allometries can take many forms and curve shapes (Mehtätalo et al., 2015). To develop our models, we considered commonly used functions (Ciceu et al., 2023). We selected three three-parameter sigmoidal functions: the Chapman – Richards (Huang et al., 1992; Richards, 1959), Hossfeld II (Elfving and Kiviste, 1997; Vallet and Perot, 2016) and Weibull (Weibull, 1951) functions.

As we aimed to develop generic models, we fitted the same equation to all the species. By comparing the Akaike Information Criterion (AIC)(Akaike, 1974), we selected the Hossfeld II function (Eq.1), which gave the best adjustments.

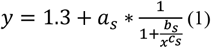

Parameters *a*_*s*_, *b*_*s*_ and *c*_*s*_ (Eq.1) were species dependent. In this function, the height equals 1.3m when the diameter tends to zero, and the sum of 1.3 + *a*_*s*_ is the asymptotic height value for large diameters. *c*_*s*_ is a shape coefficient, and *b*_*s*_ is linked by a power function to the diameter at half asymptotic height (dha) as 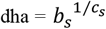. The social status of the tree was used as the explicative variable in order to take intra-plot competition into account.

#### 3.2.2 Inclusion of stand variables in relation to tree growth and biological constraints

When developing our models, we considered practical constraints related to fieldwork. Diameter measurements for all trees can be rapidly and efficiently measured, and are necessary when evaluating stand characteristics such as basal area and quadratic mean diameter. The use of a stand development stage variable, usually characterized by its dominant height, can significantly improve the fit of height-diameter allometries (Adame et al., 2008; Ciceu et al., 2023; Patrício et al., 2022). However, the quadratic mean diameter can be used as a substitute (Vallet and Perot, 2016). The quadratic diameter has the advantage of not requiring any field measurements other than tree diameter. Therefore, we chose to use basal area and quadratic mean diameter as the sole calculated stand variables, thereby simplifying the necessary field measurements.

We developed our models to quantify the influence of stand structure on height-diameter allometry. The even-aged stand structure was used as the reference condition. Other structures (uneven-aged, coppice-with-standards and coppice) were added as multiplicative adjustments representing their relative effect compared to the even-aged reference.

- *a*_*s*_ parameter

Stand variables were introduced into the asymptotic height parameter (*a*_*s*_) as formulated in equation 2:

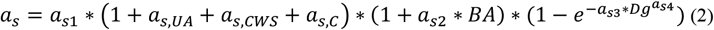

Where *a*_*s*1_ is the reference value corresponding to the even-aged structure, and *a*_*s,UA*_, *a*_*s,CWS*_ and *a*_*s,C*_ are the parameters expressing the relative difference in asymptotic height for uneven-aged, coppice-with-standards and coppice stand structures, respectively, included as dummy variables. *BA* is the basal area (in m^2^/ha) of the stand, *D*_*g*_ is the quadratic mean diameter (in cm) of the dominant species of the stand, and *a*_*s*2_, *a*_*s*3_, and *a*_*s*4_ their corresponding parameters.

The quadratic mean diameter (D_g_) was integrated into the model to reflect the biological assumption that dominant height, and therefore asymptotic height, increases as the stand ages. This effect was modeled with a negative exponential form. This form represents the relationship between quadratic diameter and dominant height, and also reflects the saturating nature of height growth over time.

The addition of basal area reflects its demonstrated influence on maximum tree height, as supported by del Río et al (2019) and Gonçalves (2022). Following similar modeling approaches (Adame et al., 2008; Temesgen and v. Gadow, 2004), we used a linear form to incorporate basal area into the asymptotic parameter. This allowed us to evaluate both the statistical significance and biological impact of stand basal area on individual tree height.

- *b*_*s*_ parameter

The *b*_*s*_ parameter, as the diameter at half asymptotic height, was developed as follows (equation 3):

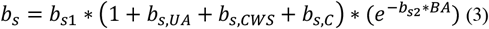

Where *b*_*s*1_ is the reference value corresponding to the even-aged structure, *b*_*s,UA*_, *b*_*s,CWS*_ and *b*_*s,C*_ are the structure-specific adjustment parameters, added as dummy variables.

Structure-specific effects were added similarly to those for *a*_*s*_. *b*_*s*2_ is the parameter associated with basal area. The exponential form for basal area was chosen to satisfy two criteria: i) the *b* parameter must remain positive to ensure realistic and biologically acceptable height estimates; ii) as basal area increases, diameter growth is limited due to competition, but height growth may continue at a similar rate (Eichhorn law, (Decourt, 1973)). Therefore, for a given diameter, small-diameter trees will be taller in a denser stand.

#### 3.2.3 Final calibrated equation

When all stand variables are taken into account, equation 4 results as follows:

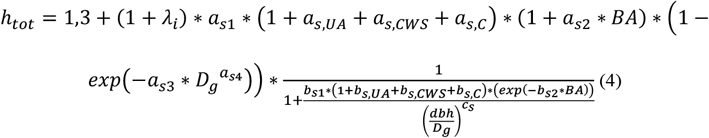

Where *h*_*tot*_ is height in m, *BA* is basal area in m^2^/ha, and *D*_*g*_ is quadratic mean diameter in cm. *λ*_*i*_ refers to the random effect calibrated (described in 3.3.1). When applying the model, its value can either be 0 (using only the fixed effects of the model) or estimated using local recalibration (described in 3.4).

### 3.3 Statistical choices

To take spatial dependency, due to trees sampled per plots, into account, we used non-linear mixed-effect models (NLME) (Pinheiro and Bates, 2006). The model included a plot random effect *λ*_*i*_, where *i* identified each plot (equation 4). The *λ*_*i*_ parameter will henceforth be referred to as the site parameter.

The residual model variance was not constant as tree height variance generally increases with tree size. Moreover, as the trees were inventoried in subplots of different radii, tree height variance was typically smaller for the smallest trees in each category, and higher for the largest. To take these two effects into account, we used a varIdent variance function from the “nlme” package (Pinheiro and Bates, 2006). We defined ten variance groups by splitting each of the five NFI diameter categories (very small trees, small trees, medium trees, large trees, and very large trees) at their median value.

### 3.4 Local recalibration process

#### 3.4.1 Principle

The local recalibration approach consisted in estimating the random site parameter (*λ*_*i*_) with data that could easily be collected in the field: the height of a small subsample of trees. This allowed the height-diameter allometric relationship to be adjusted to each plot with only a few height measurements, as illustrated in figure 1, and demonstrated in Supplementary Materials, section C.

**Figure 1:**
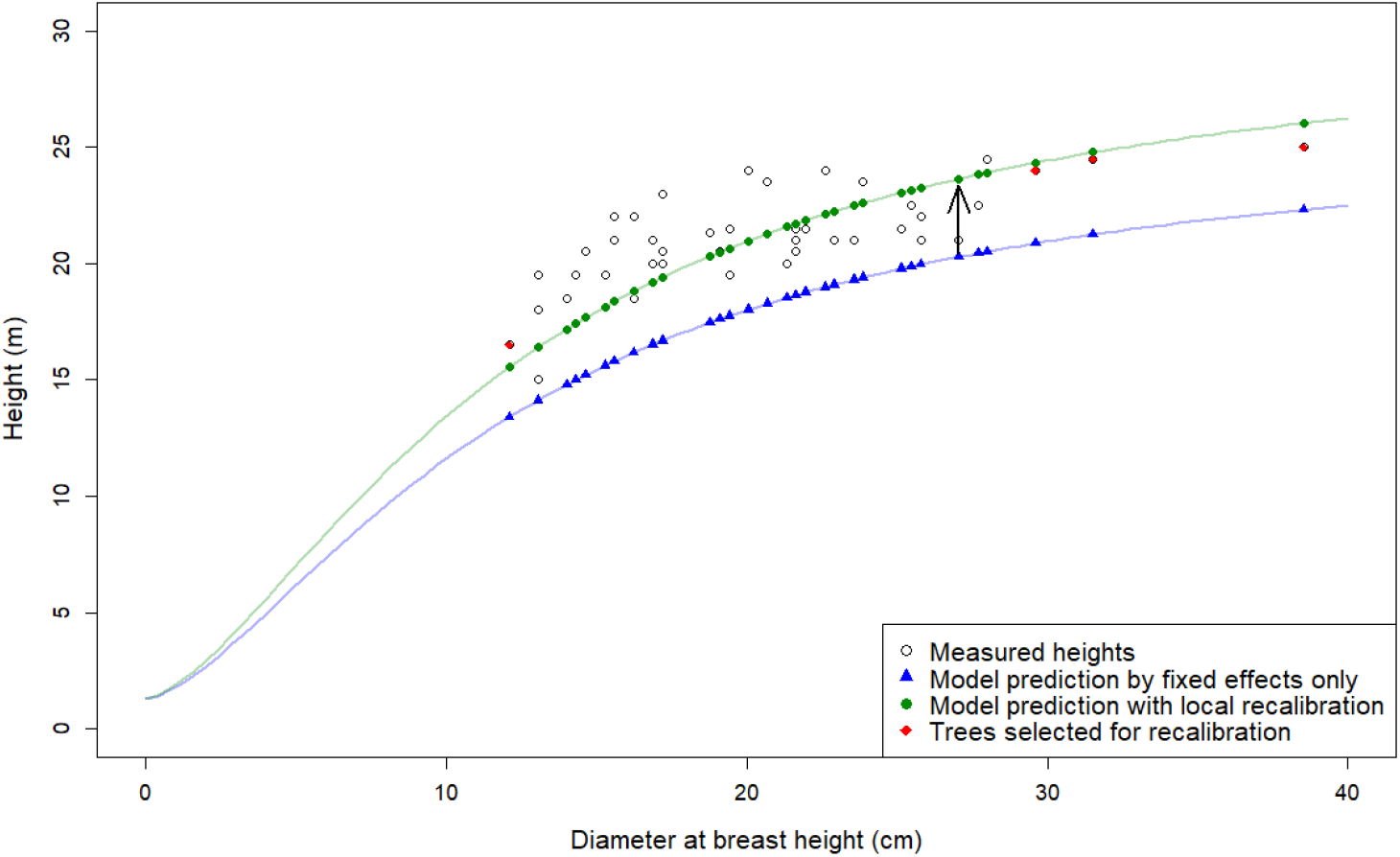
Example of local recalibration for a given Fagus sylvatica plot. The three trees with the largest diameter and the one tree with the smallest diameter were used to estimate the site parameter associated to the plot - FFA dataset.

We estimated *λ*_*m,j*_ with the mean-based approach shown in equation 5, relying only on a subset of N_j_ trees from the plot j:

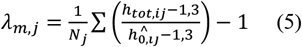

We then assessed the optimal tree subsample that would allow the site parameter to be accurately estimated and reduce the root-mean-square error (RMSE) at best, while minimizing field effort.

#### 3.4.2 Sub-samples and performance evaluation

We estimated *λ*_*m,j*_ with different samples of tree height. These samples varied in terms of size (number of trees) and in terms of relative tree diameter. We selected one to six trees according to three sampling strategies: selecting only the thinnest trees, only the largest trees, or a combination of both diameter classes. For this last strategy, we evaluated two scenarios. In the first scenario, large-diameter trees were predominant, following the distribution: 1 large; 1 large with 1 small; 2 large with 1 small; 3 large with 1 small; 3 large with 2 small and 4 large with 2 small. In the second scenario, small-diameter trees were predominant and distributed as: 1 small; 1 small with 1 large; 2 small with 1 large; 3 small with 1 large; 3 small with 2 large and 4 small with 2 large.

We applied this method to all FFA dataset plots, that had at least 13 tree records. The six largest diameter trees and the six thinnest, which were used for site parameter estimation (following the different combination described above), were then removed from the dataset. We then calculated the RMSE associated to the height prediction for each type of sub-samples, for all remaining individuals. We then identified the best diameter-class combinations based on the root-mean-square error RMSE difference relative to (i) the RMSE obtained when height was predicted using fixed effects only – and (ii) the RMSE obtained using the optimal site-parameter value, where height was predicted using *λ*_*m,j*_ as the site parameter and all available trees.

The best performing combination was then re-applied to the FFA dataset. In this second step, only the trees involved in the selected combination (ie., 6 trees of the largest, smallest or a mix of both), were removed from each plot, instead of systematically removing the six smallest and six largest diameter trees. RMSE was then calculated using all remaining individuals. To evaluate the efficiency of the local recalibration method, and identify the optimal number of trees needed, we calculated the relative RMSE improvement (RRI) as the difference between the fixed-effect model RMSE and the recalibrated model RMSE, divided by the fixed-effect RMSE.

## 4 Results

### 4.1 Calibration

Our models highlight a positive influence of quadratic mean diameter and basal area on the asymptotic height, and a negative impact of basal area on the diameter at half asymptotic height (i.e., half asymptotic height is reached for smaller diameters) (Table 3 and table S5).

**Table 3:** Model parameters and statistics for fitting of equation 4 for the ten most represented species. Significance: ***: p-value<0.001, **: p-value <0.01, *: p-value <0.05.

| Species | Estimation and Statistics |  |  |  |  |  |  |  |  |  |  |  |  |  | Residual deviation | Deviation of random effects |
| --- | --- | --- | --- | --- | --- | --- | --- | --- | --- | --- | --- | --- | --- | --- | --- | --- |
| | parameter | $a_{s1}$ | $a_{s,UA}$ | $a_{s,CWS}$ | $a_{s,C}$ | $a_{s2}$ | $a_{s3}$ | $a_{s4}$ | $b_{s1}$ | $b_{s,UA}$ | $b_{s,CWS}$ | $b_{s,C}$ | $b_{s2}$ | $c_s$ | | |
| Abies alba | estimate | 59.27 | 0.1295 | 0.0106 | - | 0.0003 | 0.0200 | 1.0637 | 0.7621 | 0.5553 | 0.0832 | - | 0.0094 | 1.5730 |  |  |
|  | significativity | *** | *** | . | - | . | *** | *** | *** | *** | *** | - | *** | *** | 2.04 | 0.13 |
|  | stddev | 2.64 | 0.01113 | 0.01003 | - | 0.00027 | 0.00108 | 0.03178 | 0.02310 | 0.02907 | 0.02282 | - | 0.00051 | 0.01615 |  |  |
| Castanea sativa | estimate | 21.36 | -0.0798 | -0.0260 | 0.0096 | 0.0031 | 0.0762 | 1.0608 | 0.3808 | 0.0305 | -0.0671 | 0.1576 | 0.0097 | 1.5726 |  |  |
|  | significativity | *** | ** | * | . | *** | *** | *** | *** | . | * | *** | *** | *** | 1.67 | 0.21 |
|  | stddev | 0.52 | 0.02736 | 0.01280 | 0.01654 | 0.00045 | 0.00926 | 0.05363 | 0.02140 | 0.08879 | 0.03017 | 0.04726 | 0.00085 | 0.03545 |  |  |
| Fagus sylvatica | estimate | 49.75 | 0.0412 | -0.0338 | -0.1867 | 0.0005 | 0.0318 | 0.9863 | 0.4410 | 0.5103 | 0.0530 | 0.1154 | 0.0029 | 1.5511 |  |  |
|  | significativity | *** | * | *** | *** | . | *** | *** | *** | *** | ** | . | *** | *** | 2.38 | 0.19 |
|  | stddev | 1.48 | 0.01789 | 0.00757 | 0.01948 | 0.00028 | 0.00134 | 0.02421 | 0.01293 | 0.05093 | 0.01709 | 0.06320 | 0.00053 | 0.01841 |  |  |
| Picea abies | estimate | 39.91 | 0.07737 | 0.01371 | - | 0.00246 | 0.01509 | 1.26384 | 0.43086 | 0.65289 | 0.06797 | - | 0.00797 | 1.93804 |  |  |
|  | significativity | *** | *** | . | - | *** | *** | . | *** | *** | * | - | *** | *** | 1.92 | 0.15 |
|  | stddev | 1.06 | 0.01273 | 0.01118 | - | 0.00028 | 0.00107 | 0.03250 | 0.01399 | 0.03865 | 0.02768 | - | 0.00049 | 0.02503 |  |  |
| Pinus pinaster | estimate | 35.06 | 0.23787 | -0.01374 | - | 0.00092 | 0.00967 | 1.40117 | 0.47802 | 1.17399 | 0.04229 | - | 0.00921 | 1.42384 |  |  |
|  | significativity | *** | *** | . | - | ** | *** | *** | *** | *** | . | - | *** | *** | 1.18 | 0.15 |
|  | stddev | 0.71 | 0.02923 | 0.00899 | - | 0.00030 | 0.00036 | 0.01761 | 0.01922 | 0.08918 | 0.02406 | - | 0.00065 | 0.02941 |  |  |
| Pinus sylvestris | estimate | 103.55 | -0.08569 | 0.01118 | - | 0.01033 | 0.00743 | 0.91605 | 0.32923 | 0.26842 | 0.04402 | - | 0.00518 | 1.81832 |  |  |
|  | significativity | . | *** | . | - | *** | . | *** | *** | *** | . | - | *** | *** | 2.32 | 0.25 |
|  | stddev | 86.87 | 0.01670 | 0.01188 | - | 0.00063 | 0.00534 | 0.06061 | 0.01348 | 0.05030 | 0.02640 | - | 0.00072 | 0.03242 |  |  |
| Pseudotsuga menziesii | estimate | 45.07 | 0.20845 | 0.01852 | - | 0.00160 | 0.00718 | 1.45561 | 0.39143 | 0.97330 | 0.05445 | - | 0.00652 | 1.74230 |  |  |
|  | significativity | *** | *** | . | - | *** | *** | *** | *** | *** | . | - | *** | *** | 1.85 | 0.13 |
|  | stddev | 1.01 | 0.0377 | 0.0111 | - | 0.0003 | 0.0004 | 0.0234 | 0.0153 | 0.1216 | 0.0299 | - | 0.0005 | 0.0338 |  |  |
| Quercus petraea | estimate | 32.63 | -0.0534 | -0.0294 | -0.0721 | 0.0060 | 0.0330 | 1.0137 | 0.1874 | 0.5170 | -0.0003 | 0.6569 | 0.0061 | 2.1660 |  |  |
|  | significativity | *** | ** | *** | *** | *** | *** | *** | *** | *** | . | *** | *** | *** | 2.02 | 0.15 |
|  | stddev | 0.54 | 0.01642 | 0.00470 | 0.01659 | 0.00032 | 0.00121 | 0.01770 | 0.00609 | 0.08860 | 0.01602 | 0.09265 | 0.00080 | 0.02605 |  |  |
| Quercus pubescens | estimate | 22.58 | -0.06542 | 0.00810 | -0.11512 | 0.01122 | 0.04416 | 0.99286 | 0.42095 | 0.06926 | 0.02891 | 0.16874 | 0.00583 | 1.70921 |  |  |
|  | significativity | *** | ** | . | *** | *** | *** | *** | *** | . | . | *** | *** | *** | 1.27 | 0.24 |
|  | stddev | 1.2 | 0.02237 | 0.01155 | 0.01390 | 0.00071 | 0.00293 | 0.04387 | 0.01796 | 0.06035 | 0.02370 | 0.04067 | 0.00090 | 0.03229 |  |  |
| Quercus robur | estimate | 30.27 | -0.08605 | 0.00685 | -0.00597 | 0.00396 | 0.03890 | 0.99302 | 0.27664 | 0.16480 | 0.02373 | 0.28099 | 0.00844 | 1.91118 |  |  |
|  | significativity | *** | *** | . | . | *** | *** | *** | *** | ** | . | *** | *** | *** | 2.11 | 0.17 |
|  | stddev | 0.55 | 0.01471 | 0.00620 | 0.02071 | 0.00033 | 0.00176 | 0.02094 | 0.00985 | 0.06260 | 0.01900 | 0.08086 | 0.00080 | 0.02957 |  |  |

The effect of stand structure on the allometric curve was species-dependent. Parameter estimates (figure 2) revealed a predominantly negative effect of coppice structure on asymptotic height compared to even-aged stands, and a strong similarity between even-aged and coppice-with-standards stands. In contrast, the effect of an uneven-aged structure on asymptotic height differed by species group, being positive for conifers and negative for broadleaved species.

**Figure 2:**
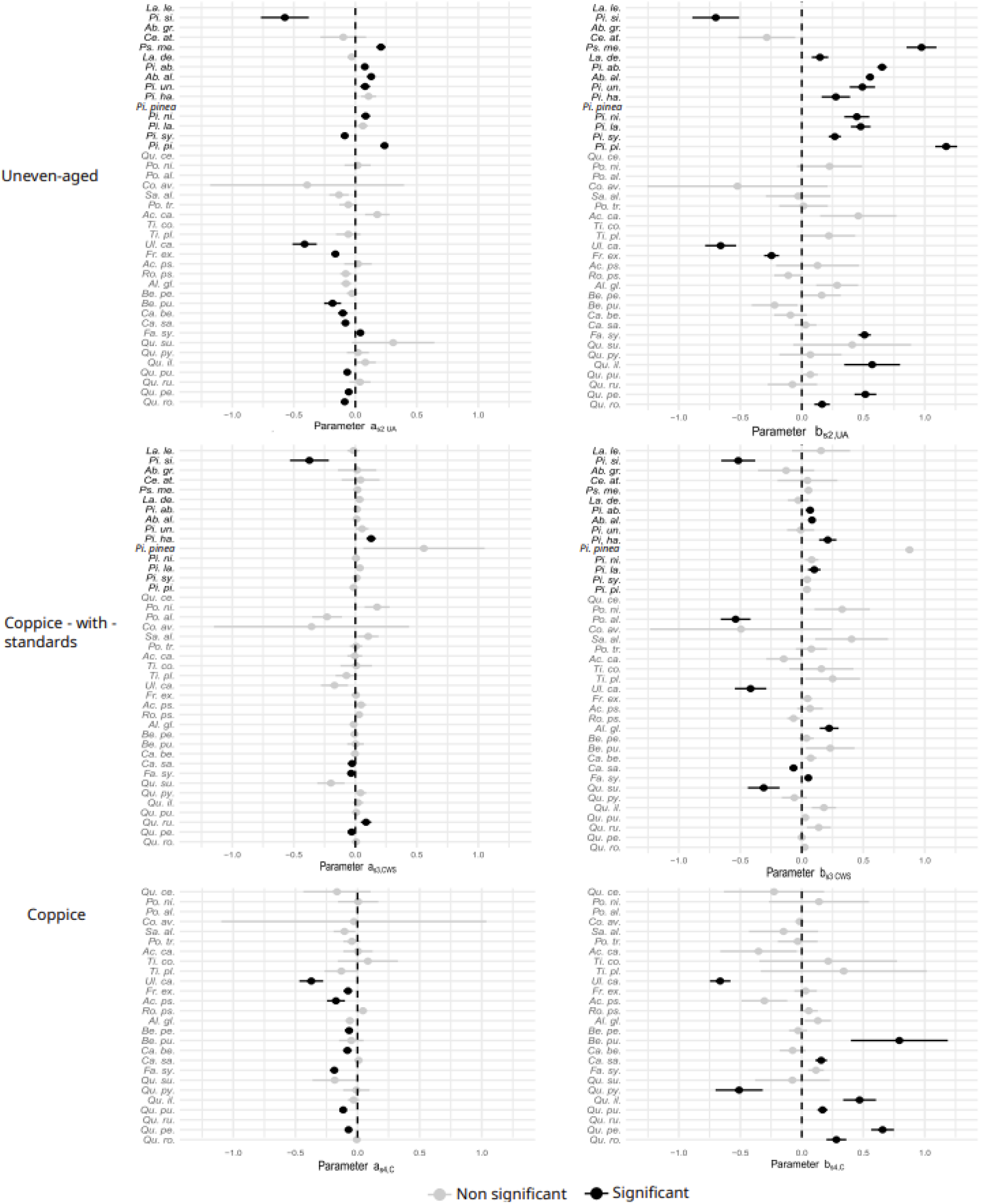
Stand structure effects on asymptotic height and diameter at half asymptotic height, compared to the even-aged reference. Estimates and associated standard deviations are presented by structure and species. Due to high non-significant values, estimates and standard deviations for Quercus cerris are not presented for parameters a_s,CWS_ and b_s,CWS_. Standard deviation was not represented for Pinus pinea for b_s,CWS_, nor for Corylus avellana for b_s,C_.

For the diameter at half asymptotic height, structural effects were substantially more mixed, with positive, negative and non-significant responses observed across species. All effects for the ten most represented species are summarized in Figure 3 and in the Supplementary Materials (figures S2 - S11).

**Figure 3:**
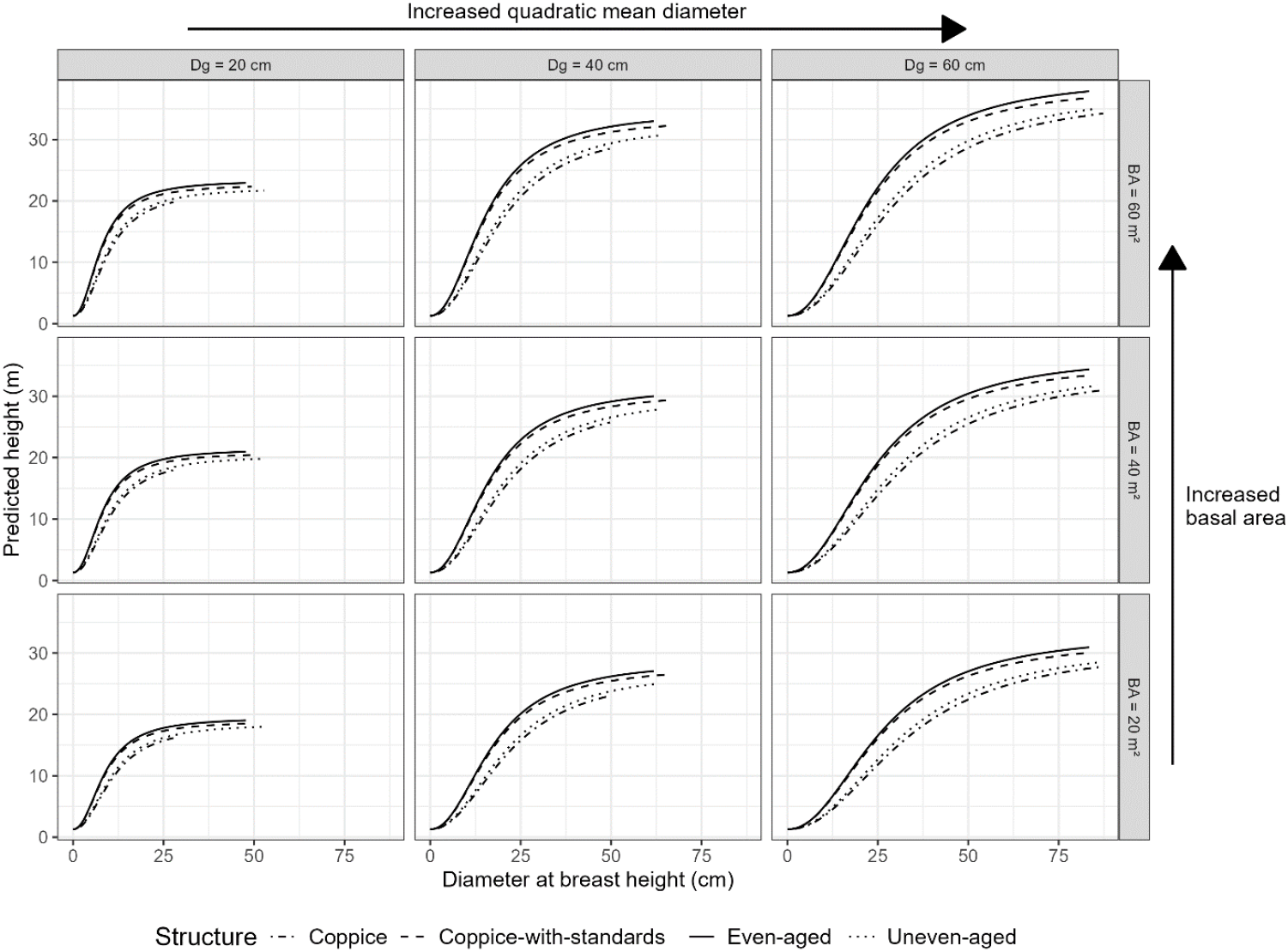
Presentation of the Quercus petraea model, for varying basal area, quadratic diameter and stand characteristics. Predictions are limited to the range of diameter values observed for each structure in the calibration dataset.

### 4.2 Validation (fixed effects only)

We calculated the errors associated to *ĥ*_*0*_. Without the site parameter, height estimates were contained between three and five meters for the calibration dataset. Table 4 highlights similar results for model errors between the calibration and NFI_2023 datasets. As expected, the results for the FFA dataset were less similar to the calibration dataset. A bias in estimation can be observed when predicted height is confronted to observed height (figure 4). Models’ residual graphs are presented in supplementary materials, section G. Results of the one-sample t-test showed a significant bias of -0.22m on the NFI_2023 dataset. For the FFA dataset, the t-test showed a significant difference of 2.75m.

**Table 4:** Validation results for 6 species across all datasets. For the calibration dataset and NFI_2023, results aggregate all stand structure types. Only even-aged structures are present in the FFA dataset.

| Species | Calibration dataset |  | NFI_2023 |  | FFA |  |
| --- | --- | --- | --- | --- | --- | --- |
|  | RMSE<sub>0</sub>(m) | Nb obs. | RMSE<sub>0</sub>(m) | Nb obs. | RMSE<sub>0</sub>(m) | Nb obs. |
| *Abies alba* | 3.5551 | 19 055 | 3.5644 | 606 | 2.7906 | 618 |
| *Fagus sylvatica* | 4.535 | 28 135 | 5.1654 | 853 | 5.0553 | 816 |
| *Picea abies* | 3.7235 | 19 176 | 3.8088 | 390 | 4.4948 | 287 |
| *Pinus sylvestris* | 3.8592 | 21 865 | 4.2631 | 496 | 5.3985 | 246 |
| *Pseudotsuga menziesii* | 3.3942 | 13 691 | 3.3494 | 475 | 2.611 | 212 |
| *Quercus petraea* | 3.5309 | 31 927 | 3.6339 | 1 187 | 3.5484 | 830 |

**Figure 4:**
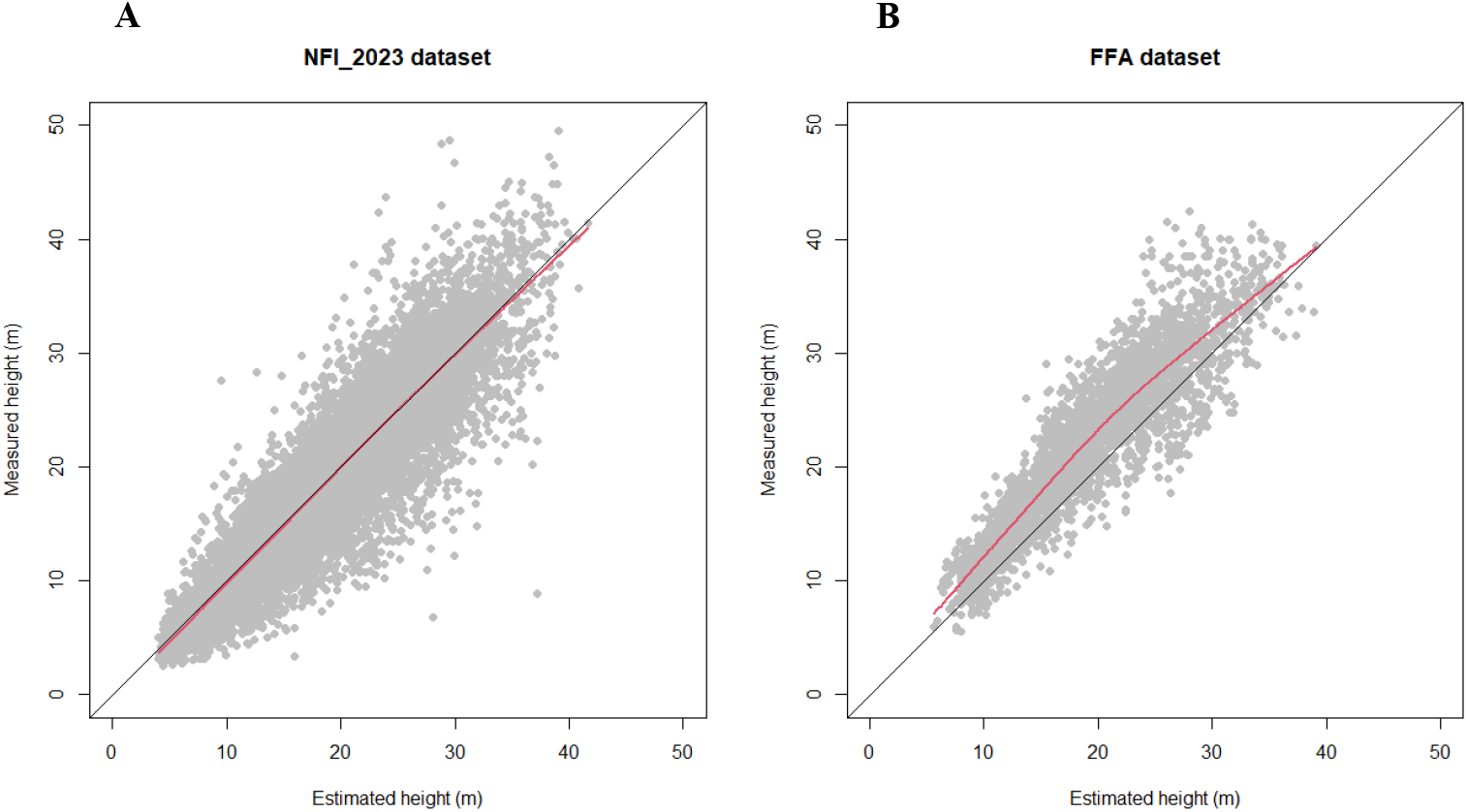
Observed height compared to predicted height for the whole NFI_2023 dataset (A – 40 species; 8771 observations) and the FFA dataset (B – 6 species; 3009 observations). The black line represents perfect prediction (1:1). The red line shows the overall trend in the data (lowess regression).

### 4.3 Local recalibration

The selection of a combination of trees with a predominance in trees with the largest diameter reduced RMSE the most (figure 5). From one to three trees, adding an additional tree provided a strong reduction in RMSE; thereafter, adding more trees only slightly improved the reduction in RMSE.

**Figure 5:**
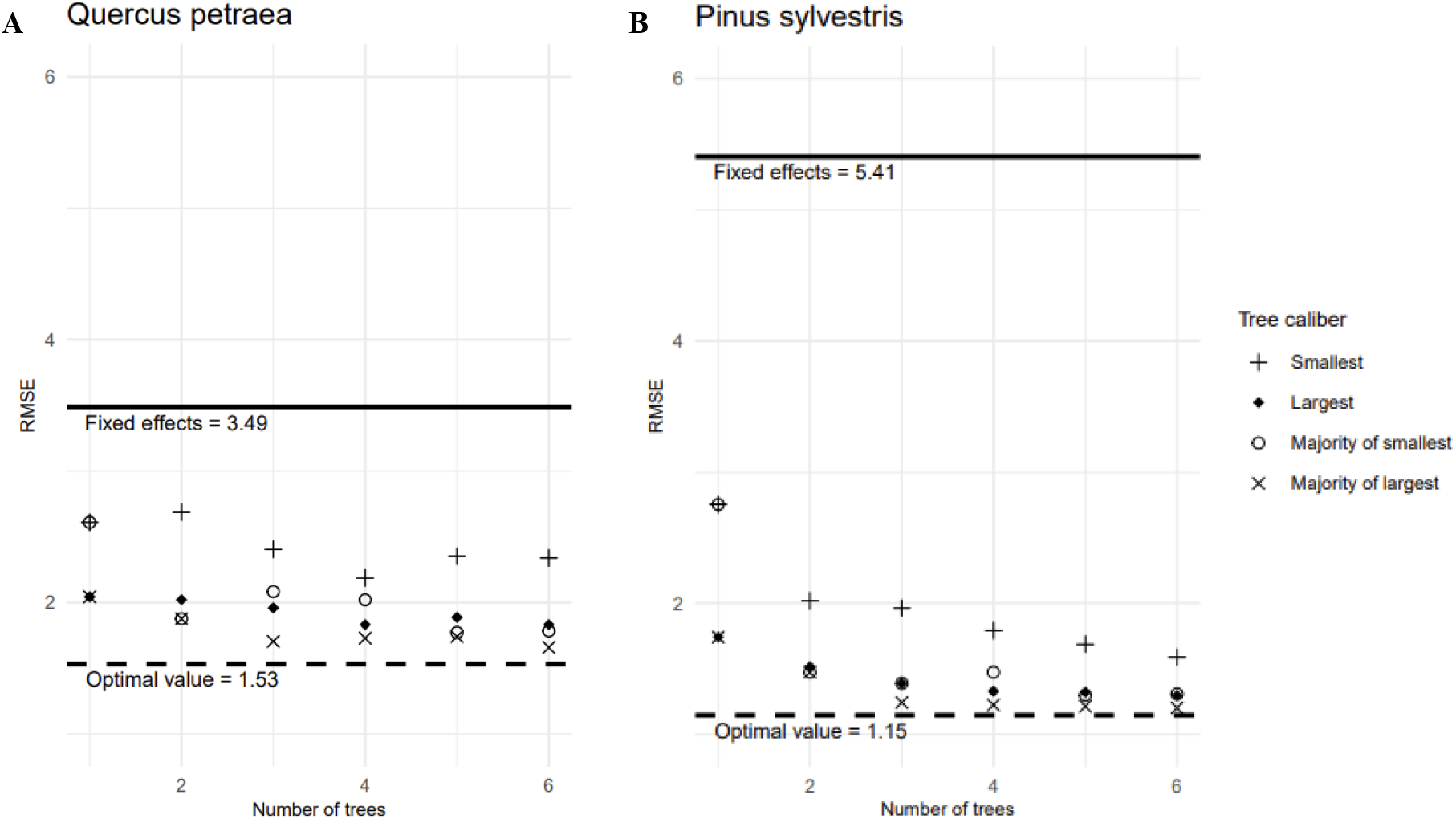
RMSE values for tree height prediction with recalibration, with different tree sub-samples based on number and diameter class of the trees on the calibration dataset, per species (Quercus petraea (A) and Pinus sylvestris (B))

The root-mean-square error of tree height predictions following local recalibration applied to the FFA dataset, with sub-samples of one to six trees with a predominance of large diameter trees, are presented in table 5. Estimating the site parameter with all trees present on the plot (optimal value) consistently improved prediction accuracy across species, with relative RMSE improvement (RRI) ranging from 20 to 70 %, compared to fixed effects.

**Table 5:** Local recalibration results for the FFA dataset, compared to fixed-effects-only height prediction. RMSE in meters and relative RMSE improvement (RRI) in percentage, for height predictions with fixed effects, optimal RMSE value, and sub-samples of one to six trees selected among the largest and smallest diameters with predominant large trees, for the FFA dataset. RRI is calculated relative to the fixed-effects baseline.

| Site parameter estimates |  | Abies alba |  | Fagus sylvatica |  | Picea abies |  | Pinus sylvestris |  | Pseudotsuga menziesii |  | Quercus petraea |  |
| --- | --- | --- | --- | --- | --- | --- | --- | --- | --- | --- | --- | --- | --- |
|  |  | RMSE (m) | RRI (%) | RMSE (m) | RRI (%) | RMSE (m) | RRI (%) | RMSE (m) | RRI (%) | RMSE (m) | RRI (%) | RMSE (m) | RRI (%) |
|  | Fixed effects | 2.66 | - | 5.06 | - | 4.40 | - | 5.46 | - | 2.65 | - | 3.51 | - |
|  | Optimal value | 1.98 | 25.76 | 1.84 | 63.60 | 2.42 | 44.86 | 1.21 | 77.77 | 1.26 | 52.39 | 1.57 | 55.45 |
| Number of trees selected among the largest | 1 | 3.00 | -12.64 | 3.44 | 31.92 | 2.87 | 34.82 | 1.90 | 65.21 | 2.57 | 2.82 | 2.15 | 38.94 |
|  | 2 | 3.35 | -25.56 | 2.29 | 54.79 | 3.70 | 15.95 | 1.53 | 72.02 | 2.81 | -6.04 | 1.89 | 46.33 |
|  | 3 | 2.82 | -5.66 | 2.06 | 59.38 | 3.08 | 29.95 | 1.33 | 75.68 | 2.41 | 8.81 | 1.76 | 49.95 |
|  | 4 | 2.69 | -0.85 | 2.15 | 57.57 | 2.80 | 36.27 | 1.31 | 75.97 | 2.22 | 16.08 | 1.77 | 49.57 |
|  | 5 | 2.50 | 6.29 | 2.02 | 60.04 | 2.95 | 32.92 | 1.30 | 76.24 | 2.17 | 17.85 | 1.76 | 49.99 |
|  | 6 | 2.42 | 9.06 | 2.05 | 59.52 | 2.74 | 37.75 | 1.28 | 76.51 | 1.99 | 24.96 | 1.69 | 52.03 |
| Number of trees |  | 480 |  | 652 |  | 245 |  | 176 |  | 176 |  | 648 |  |

When the site parameter was estimated from a sub-sample of three to four trees (2 large with 1small and 3large with 1 small), substantial improvements in RMSE generally resulted, compared to using fixed effects only. The RRI ranges from 10 to 70%, except for *Abies alba*, for which the use of sub-samples led to higher RMSEs than the fixed-effects baseline. This is likely due to *Abies alba* plots displaying a height-diameter relationship closer to those of the calibration dataset than those of the other species.

## 5. Discussion

### 5.1 Effects of stand characteristics on height-diameter allometry

#### 5.1.1 Influence of stand structure

Stand structure had a significant effect on height-diameter allometry for 30 species. However, a few species, such as *Populus tremula*, showed no significant parameter estimates for any of the stand structures. Overall, the stand structure parameter estimates (figure 2) highlight the influence of forest management on allometric relationships.

In coppice stands, results were consistent with the regeneration method (vegetative sprouting from existing root system), which generally leads to reduced tree height growth compared to even-aged stands for a given diameter. However, the diameter at half asymptotic height (dha) was often greater than in even-aged conditions. This shift suggests a reallocation of resources toward radial rather than vertical growth, a pattern observed in previous studies (Corcuera et al., 2006; Deforce and Haneca, 2015).

In uneven-aged stands, competition between individuals is generally reduced compared to that in even-aged stands, because density is lower. As highlighted by del Rìo et al. 2019, higher competition leads to greater tree heights for a given diameter, due to the increased drive for vertical growth under denser conditions. This pattern is consistent with the fact that reduced competition, particularly for light in uneven-aged stands, allows individuals to allocate more resources to radial growth than to height. As a result, trees in uneven-aged stands tend to be of smaller height for a given diameter, leading to a negative effect on asymptotic height compared to trees in even-aged stands. Our results for broadleaved species were consistent with this observation. However, coniferous species showed a positive effect on asymptotic height, which can partly be explained by the correlation between asymptotic height and dha parameters: the diameter range for these species corresponds to a section of the curves where height is greater for even-aged stands even if the asymptotic height is lower (the same effect as in Supplementary Materials, section F).

Additionally, our results indicate an increase in dha in uneven-aged stands, which is consistent with lesser light competition compared to even-aged stands. Under high competition, such as in even-aged stands, trees prioritize height growth over radial growth, shifting the allometry curve to the left. Therefore, for a given diameter (among small trees), trees in uneven-aged stands tend to be shorter than those in even-aged stands (trends for parameters *a* and *b* in the HossfledII equation (Vallet and Perot, 2016)).

#### 5.1.2 Influence of basal area

Tree height for a given diameter can increase with stand density. Basal area (BA), often used as a proxy for stand density, also serves as an index for competition among trees. As basal area increases, competition for resources intensifies, and trees tend to allocate more energy to vertical growth in order to maintain access to light. Consequently, for a given diameter, tree height is expected to be greater in denser stands, as highlighted by del Río et al. (2019) and Gonçalves (2022).

In our models, this effect is reflected through an increase in asymptotic height and a corresponding decrease in the diameter at half asymptotic height (parameters *a*_*s*2_ and *b*_*s*2_), consistent with our expectations. As a result, the height-diameter allometric curve shifts upward: for a given diameter, predicted height increases with basal area (Figure 2). However, this trend did not apply uniformly across all species. For *Corylus avellana, Pinus halepensis* and *Tilia cordata*, basal area had a significant negative effect on asymptotic tree height. As illustrated in figure S12 (Supplementary Materials, section F), in the case of *Tilia cordata*, the high effect of basal area over diameter at half asymptotic height, associated with a lack of observation for large diameters, tended to bend the allometric curve to the left, inducing a reduction in asymptotic height by mathematical construction. Hence, as for some effects in uneven-aged stands, for the observed range of diameters of these species, the height is still higher with increasing BA even if the asymptotic parameter is negative.

In addition to its effect on asymptotic height, increased competition directly influences diameter growth. As emphasized by Aleinikovas et al. (2014), increased competition has a negative effect on tree diameter growth. As a consequence, the height-diameter curve is shifted to the left, indicating greater heights for a given diameter.

#### 5.1.3 Influence of quadratic mean diameter

Quadratic mean diameter provides insights into the developmental stage of the stand (Aussenac et al., 2023; Sharma et al., 2019a). The silvicultural system applied to each structure will induce different growth dynamics. For example, even-aged stands are typically allowed to grow for several decades, enabling trees to reach greater heights and diameters. In contrast, coppice systems imply periodic cutting, and the trees regenerate from stump sprouts, resulting in shorter growth periods, and more limited development in size. In both uneven-aged and coppice structures, Dg values tended to remain relatively stable over time, due to the continuous or cyclical nature of their management regimes.

Across all species studied, Dg had a positive effect on the asymptotic height in the height diameter allometry. As Dg increased, the height asymptote also increased. This pattern aligns with findings in the literature, which report a strong relationship between Dg, dominant height, and height-diameter allometry (Schmidt et al., 2018; Sharma et al., 2019a). The increase in asymptotic height with Dg reflects a typical stand development trajectory, where older and more developed stands support taller dominant trees.

### 5.2 Optimization of model accuracy and predictive performance

#### 5.2.1 Quality of the models

Using National Forest Inventory (NFI) data enabled the inclusion of a wide range of species and tree records, covering diverse climatic, structural and edaphic conditions across the French mainland territory. The robustness of the dataset increases the confidence in applying our models across Europe as well, as was proven possible for studies on growth (Mahnken et al., 2022). Additionally, the large size of the dataset reduced the influence of outliers during calibration, enhancing parameter stability and reliability. The RMSE obtained for each species are in line with other studies, where the models selected had a RMSE between 1 and 3.5 meters (Adame et al., 2008; Gómez-García et al., 2015; Mehtätalo et al., 2015; Vázquez-Veloso et al., 2025).

To assess the efficiency of including a stand structure effect in our models, we evaluated the RMSE associated with applying the even-aged calibrated model to plots with different stand structures and compared it to structure-specific model errors. Estimations were done using only fixed effects. The relative error difference calculated for the calibration dataset (Table 6) indicates a strong improvement in model accuracy with structure-specific models for uneven-aged and coppice stand structures. In contrast, applying the even-aged model to coppice-with-standards plots revealed a high degree of similarity between the two structures, as reflected by the relatively small RMSE differences. As mentioned in Gonçalves (2022), using even-aged models to predict tree height in uneven-aged stands can lead to estimation errors of up to 20%. Our models show similar degrees of difference between estimation errors (table 6). Our results were obtained with only the fixed effects of our models, which explains the negative result for *Pseudotsuga menziesii*. Thus, adding stand structure to our models improved the accuracy of estimated tree height across diverse management systems, and provided insights into less studied structures and how they differ from the even-aged structure.

**Table 6:** Relative difference in RMSE (%) when applying models calibrated for even-aged plots to stands with other structures. Positive values indicate higher prediction error for the even-aged model than for structure-specific models, while negative values indicate better accuracy for the even-aged model. These results are based on the calibration dataset.

| Species | Even-aged model applied to stand structure |  |  |
| --- | --- | --- | --- |
|  | Uneven-aged | Coppice-with-standards | Coppice |
| *Abies alba* | +6.8 | +1.2 |  |
| *Castanea sativa* | +15.5 | +1.9 | +2.0 |
| *Fagus sylvatica* | +11.7 | +5.3 | +50.8 |
| *Picea abies* | +8.4 | +0.2 |  |
| *Pinus pinaster* | +9.8 | +3.6 |  |
| *Pinus sylvestris* | +26.4 | -0.3 |  |
| *Pseudotsuga menziesii* | -1.2 | -0.3 |  |
| *Quercus petraea* | +16.5 | +3.9 | +24.8 |
| *Quercus pubescens* | +10.9 | -0.2 | +21.8 |
| *Quercus robur* | +19.5 | -0.2 | +6.4 |

Finally, model bias observed on validation datasets can largely be attributed to the lack of site-specific information. This limitation arises from applying a generalized model to a specific region, a known source of error when site fertility or climate diverges from the average conditions of the calibration dataset (Mensah et al., 2018). Nevertheless, the low bias observed on the NFI_2023 dataset suggests high consistency with the calibration data. In contrast, the underestimation of tree height in the FFA dataset was expected, as these plots concern more fertile areas than do the calibration plots.

#### 5.2.2 Benefits of local recalibration

This approach significantly improved performance on the FFA dataset, especially where site fertility deviated from the average. Our recalibration strategy focused on a predominant number of trees with the largest diameters, in line with recommendations from Sullivan et al. (2018), who showed that using large-diameter trees improves recalibration outcomes. Furthermore, as shown by Sharma and Breidenbach (2015), including trees from different species when estimating site parameters induces error. We therefore performed species-specific recalibrations.

### 5.3 Limits

We chose to develop a single model structure that could be adapted to different stand types by adding structure parameters. While this approach simplifies generalization and cross-species comparison, it may have limitations. For example, coppice systems, which are managed through frequent cutting cycles, pose particular modeling challenges and can complicate convergence when the same stand variables as those applied to other structures are used. In uneven-aged stands, the management process tends to maintain basal area and quadratic mean diameter within a narrow range, unlike in even-aged stands. Consequently, using the same variables to characterize structurally distinct stands may reduce model relevance or performance in some cases.

We also used a single functional form across species, selected based on the overall AIC comparison. For a few less-represented species, alternative functional forms would have produced lower AIC values. Developing species-specific functions could improve model accuracy, but maintaining a consistent equation enables parameter comparisons across species and the development of generic models for practical use. We also used species-specific models to test each species’ response to competition and stand dynamics.

### 5.4 Implications for forest modelling and management

The models developed in this study account for both stand structure and dendrometrical characteristics, providing highly accurate height estimates across different management systems (even-aged, uneven-aged, coppice and coppice-with-standards). This represents a notable advancement in modeling, and can improve estimates of other critical forest variables such as timber volume and carbon stocks.

Our validation results show strong predictive performance in plots with characteristics similar to the calibration dataset. In plots with high fertility or atypical conditions, applying local recalibration reduced bias (see practical method in Supplementary Materials, section C). The number of trees required for recalibration can be adjusted depending on the desired accuracy of the height estimates, but as few as one to four trees could already strongly improve the results (Table 5).

## 6 Conclusion

Based on data from more than 40,000 stands surveyed by the French National Forest Inventory, we developed height-diameter allometries for 41 European species that take into account stand characteristics.

We found that stand structure (even-aged, uneven-aged, coppice-with-standards and coppice) did significantly impact height-diameter allometry, as did the quadratic mean diameter and the basal area of the stand. Given the large diversity of climates, stand structures and soil fertility present in the French metropolitan area, the models developed here could be applied all over Europe. Moreover, to address the loss of local accuracy when applying a large-scale model at stand level, we developed a recalibration method requiring only a few height measurements on the largest trees in the stand. Measuring one to four trees can considerably reduce estimation errors.

## Declarations

### Availability of data and material

Data from the French National Forest Inventory are available online.

## Author’s contributions

Natacha Savine: Writing – original draft, Validation, Methodology, Investigation, Data curation, Conceptualization.

Thomas Cordonnier: Writing – review & editing, Supervision, Methodology, Investigation, Conceptualization.

Gauthier Ligot: Writing – review & editing, Supervision, Methodology, Investigation.

Patrick Vallet: Writing – review & editing, Supervision, Methodology, Investigation, Funding acquisition, Conceptualization.

## Acknowledgments

The authors wish to thank the French National Forest Office, who funded this work under the grant “Régénération, croissance et production des forêts de plaine, 2022-2024”. The authors also thank the National Forest Inventory for making the data available free of charge.

