## supplementary materials for "Robust height-diameter allometries for 41 European tree species: stand characteristics and structure matter"

### A. Description of stand structures

*Figure S 1: Diversity of stand structure in the NFI dataset throughout the French metropolitan territory (2022)*

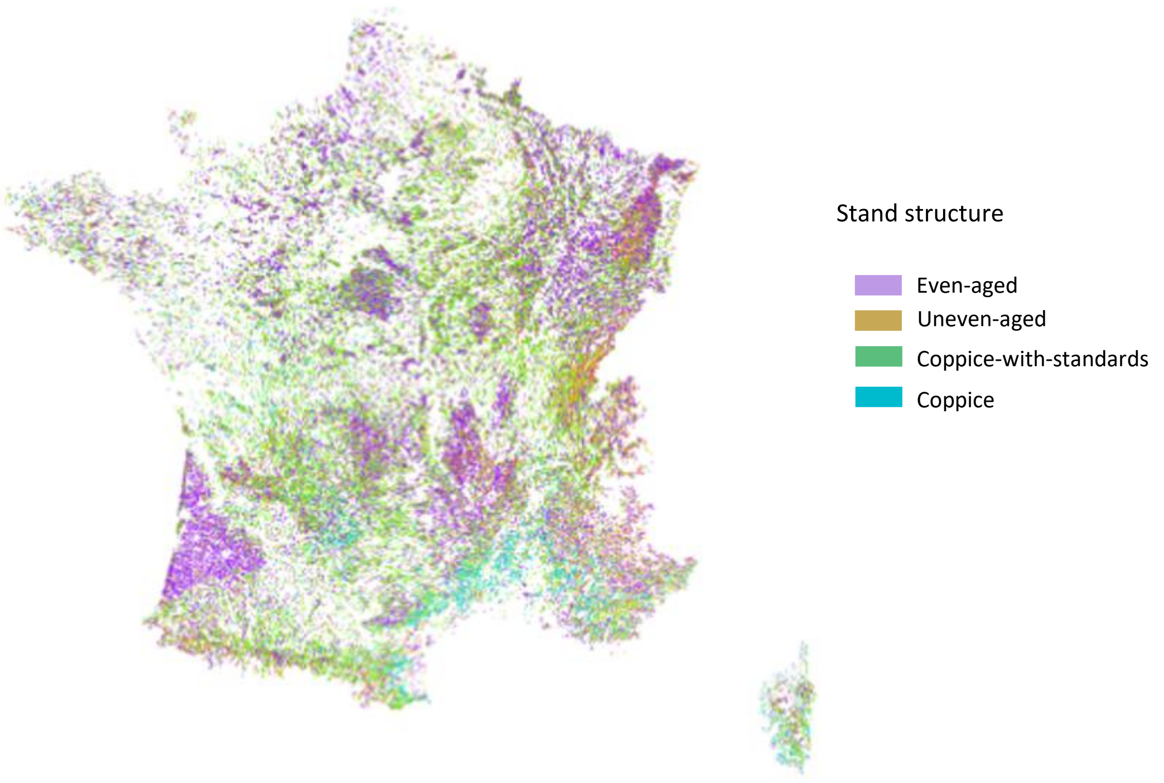


*Table S 1: Correspondences between structure variables*

| Sfo (2005 - 2013) | Sver (2014 - present ) |
|---|---|
| Even-aged | Regular (closed) stand with a low canopy layer / High regular structure without understory |
| Uneven-aged | Irregular vertical structure |
| Coppice-with-standards | High regular structure with understory |
| Coppice | Other low regular stand |
| Neither structure nor information | Neither structure nor information |

### B. Description of the study datasets

*Table S 2: Description of the 41 species in the calibration dataset. EA: Even-aged, UA: Uneven-aged, CWS: Coppice-With-Standards, C: Coppice*

| Species | Number of obs. | Number of plots | Number of obs. per structure | | | | DBH (cm) | | | Height (m) | | |
|---|---|---|---|---|---|---|---|---|---|---|---|---|
| | | | EA | UA | CWS | C | min | mean | max | min | mean | max |
| *Abies alba* | 19,055 | 2,361 | 12,502 | 3,772 | 2,781 | 0 | 7.48 | 36.91 | 129.23 | 2.4 | 21.90 | 46.2 |
| *Abies grandis* | 306 | 35 | 262 | 0 | 44 | 0 | 7.64 | 29.95 | 61.18 | 3.9 | 20.31 | 37.0 |
| *Acer campestre* | 118 | 55 | 15 | 13 | 51 | 39 | 7.64 | 16.52 | 56.66 | 3.1 | 7.99 | 16.2 |
| *Acer pseudoplatanus* | 845 | 212 | 358 | 46 | 367 | 74 | 7.54 | 24.95 | 92.69 | 4.1 | 17.33 | 42.9 |
| *Alnus glutinosa* | 2,690 | 501 | 816 | 176 | 1,247 | 451 | 7.48 | 25.10 | 136.87 | 3.3 | 17.66 | 37.5 |
| *Betula pendula* | 2,730 | 654 | 757 | 143 | 1,206 | 624 | 7.48 | 17.98 | 68.12 | 2.0 | 14.93 | 34.1 |
| *Betula pubescens* | 253 | 66 | 74 | 37 | 94 | 48 | 7.54 | 20.08 | 65.67 | 4.3 | 15.55 | 28.9 |
| *Carpinus betulus* | 4,536 | 1,105 | 1,698 | 148 | 2,390 | 300 | 7.51 | 21.96 | 100.01 | 3.2 | 17.00 | 36.7 |
| *Castanea sativa* | 14,334 | 2,670 | 2,993 | 417 | 7,050 | 3,874 | 7.48 | 27.46 | 262.92 | 2.2 | 15.40 | 34.8 |
| *Cedrus atlantica* | 297 | 73 | 218 | 16 | 63 | 0 | 7.64 | 22.10 | 117.77 | 3.3 | 11.74 | 33.8 |
| *Corylus avellana* | 263 | 120 | 18 | 64 | 106 | 75 | 7.48 | 10.81 | 30.24 | 3.9 | 8.89 | 14.8 |
| *Fagus sylvatica* | 28,135 | 4,402 | 16,437 | 1,302 | 9,201 | 1,195 | 7.48 | 33.52 | 174.12 | 1.3 | 20.21 | 47.1 |
| *Fraxinus excelsior* | 5,523 | 1,139 | 2,36 | 375 | 2,381 | 407 | 7.48 | 25.41 | 120.80 | 2.5 | 18.84 | 40.0 |
| *Larix decidua* | 2,703 | 420 | 1,925 | 531 | 247 | 0 | 7.54 | 33.54 | 151.52 | 3.1 | 18.25 | 42.0 |
| *Larix kaempferi* | 245 | 55 | 193 | 0 | 52 | 0 | 8.59 | 28.38 | 53.51 | 6.0 | 21.09 | 36.5 |
| *Picea abies* | 19,176 | 2,664 | 14,933 | 2,127 | 2,116 | 0 | 7.51 | 32.14 | 127.01 | 1.6 | 21.68 | 48.4 |
| *Picea sitchensis* | 1,562 | 207 | 1,356 | 14 | 192 | 0 | 7.64 | 33.79 | 109.66 | 3.2 | 21.24 | 39.1 |
| *Pinus halepensis* | 4,216 | 849 | 2,336 | 425 | 1,455 | 0 | 7.54 | 28.95 | 99.47 | 3.0 | 12.37 | 27.3 |
| *Pinus laricio* | 5,116 | 857 | 3,959 | 354 | 803 | 0 | 7.48 | 29.29 | 133.37 | 2.9 | 16.31 | 45.4 |
| *Pinus nigra nigra* | 4,353 | 763 | 3,050 | 359 | 944 | 0 | 7.64 | 26.03 | 79.58 | 2.8 | 14.28 | 37.0 |
| *Pinus pinaster* | 20,695 | 4,075 | 16,204 | 551 | 3,940 | 0 | 7.48 | 30.61 | 111.09 | 2.6 | 16.93 | 39.1 |
| *Pinus pinea* | 313 | 62 | 212 | 0 | 101 | 0 | 7.58 | 25.89 | 74.80 | 3.1 | 8.82 | 20.6 |
| *Pinus sylvestris* | 21,865 | 3,572 | 15,409 | 1,988 | 4,468 | 0 | 7.48 | 27.22 | 98.96 | 1.7 | 13.97 | 41.2 |
| *Pinus uncinata* | 1,900 | 272 | 878 | 790 | 232 | 0 | 7.64 | 26.37 | 81.30 | 2.4 | 10.53 | 22.5 |
| *Populus alba* | 113 | 13 | 47 | 0 | 66 | 0 | 7.96 | 44.98 | 117.77 | 7.0 | 22.96 | 41.0 |
| *Populus nigra* | 399 | 87 | 140 | 80 | 144 | 35 | 7.64 | 39.59 | 158.49 | 4.0 | 20.95 | 43.5 |
| *Populus tremula* | 1,038 | 238 | 294 | 78 | 440 | 226 | 7.48 | 20.05 | 87.22 | 3.1 | 16.72 | 35.6 |
| *Pseudotsuga menziesii* | 13,691 | 2,151 | 11,809 | 207 | 1,675 | 0 | 7.51 | 33.20 | 140.69 | 2.0 | 23.62 | 49.4 |
| *Quercus cerris* | 125 | 23 | 81 | 0 | 24 | 20 | 7.64 | 32.87 | 79.90 | 5.2 | 19.92 | 35.4 |
| *Quercus ilex* | 5,150 | 1,756 | 393 | 231 | 1,372 | 3,154 | 7.48 | 17.28 | 198.94 | 1.7 | 7.56 | 24.5 |
| *Quercus petraea* | 31,927 | 5,772 | 16,779 | 575 | 13,370 | 1,203 | 7.48 | 35.64 | 137.35 | 1.9 | 20.95 | 44.0 |
| *Quercus pubescens* | 20,689 | 4,655 | 5,527 | 879 | 9,188 | 5,095 | 7.48 | 20.80 | 192.26 | 1.8 | 11.39 | 30.8 |
| *Quercus pyrenaica* | 510 | 146 | 231 | 53 | 192 | 34 | 7.48 | 25.01 | 85.63 | 4.0 | 13.99 | 27.0 |
| *Quercus robur* | 28,049 | 5,691 | 13,362 | 935 | 12,762 | 990 | 7.48 | 35.68 | 164.12 | 2.2 | 18.83 | 43.2 |
| *Quercus rubra* | 896 | 183 | 674 | 31 | 191 | 0 | 7.54 | 23.89 | 115.23 | 4.5 | 18.08 | 40.6 |
| *Quercus suber* | 984 | 211 | 355 | 57 | 472 | 100 | 7.64 | 30.58 | 117.14 | 2.8 | 7.88 | 20.8 |
| *Robinia pseudoacacia* | 3,301 | 674 | 666 | 117 | 1,728 | 790 | 7.51 | 18.70 | 122.33 | 5.0 | 15.82 | 37.6 |
| *Salix alba* | 375 | 89 | 93 | 51 | 160 | 71 | 7.64 | 38.90 | 131.78 | 3.1 | 17.30 | 32.9 |
| *Tilia cordata* | 243 | 49 | 69 | 0 | 146 | 28 | 7.64 | 24.44 | 82.51 | 3.3 | 16.94 | 30.0 |
| *Tilia platyphyllos* | 238 | 45 | 56 | 36 | 121 | 25 | 7.64 | 25.21 | 101.22 | 5.7 | 15.55 | 29.0 |
| *Ulmus minor* | 237 | 71 | 52 | 23 | 63 | 99 | 7.51 | 13.14 | 41.06 | 3.5 | 9.57 | 19.4 |

*Table S 3: Species in the NFI_2023 dataset*

| Species | Number of obs | Number of plots | Number of obs. per structure | | | | dbh (cm) | | | Height (m) | | |
|---|---|---|---|---|---|---|---|---|---|---|---|---|
| | | | EA | UA | CWS | C | min | mean | max | min | mean | max |
| Abies alba | 606 | 120 | 276 | 95 | 235 | 0 | 7.61 | 42.18 | 104.02 | 3.9 | 23.86 | 44.8 |
| Acer campestre | 19 | 8 | 8 | 4 | 7 | 0 | 7.51 | 15.63 | 30.37 | 7.1 | 12.78 | 18.6 |
| Acer pseudoplatanus | 71 | 25 | 26 | 8 | 32 | 5 | 7.86 | 25.08 | 64.49 | 5.8 | 17.53 | 31.5 |
| Alnus glutinosa | 93 | 30 | 19 | 21 | 53 | 0 | 7.51 | 25.50 | 59.59 | 7.5 | 18.16 | 30.0 |
| Betula pendula | 54 | 24 | 16 | 3 | 32 | 3 | 7.70 | 15.80 | 41.48 | 5.2 | 12.71 | 22.9 |
| Betula pubescens | 37 | 11 | 10 | 10 | 17 | 0 | 7.64 | 22.10 | 40.78 | 8.2 | 16.46 | 26.3 |
| Carpinus betulus | 147 | 51 | 39 | 11 | 95 | 2 | 7.58 | 22.40 | 56.91 | 7.5 | 17.94 | 30.6 |
| Castanea sativa | 365 | 110 | 76 | 45 | 238 | 6 | 7.48 | 34.86 | 229.82 | 4.3 | 15.66 | 33.1 |
| Cedrus atlantica | 13 | 5 | 9 | 2 | 2 | 0 | 8.37 | 20.10 | 29.41 | 6.1 | 11.45 | 16.7 |
| Corylus avellana | 10 | 6 | 0 | 5 | 5 | 0 | 7.58 | 10.94 | 13.91 | 7.5 | 9.91 | 12.3 |
| Fagus sylvatica | 853 | 210 | 335 | 48 | 467 | 3 | 7.54 | 40.42 | 136.24 | 5.3 | 22.98 | 48.7 |
| Fraxinus excelsior | 140 | 47 | 26 | 34 | 80 | 0 | 7.80 | 27.97 | 74.80 | 5.6 | 19.00 | 41.4 |
| Larix decidua | 76 | 21 | 55 | 6 | 15 | 0 | 7.54 | 44.59 | 105.46 | 5.4 | 21.20 | 35.8 |
| Larix kaempferi | 10 | 2 | 10 | 0 | 0 | 0 | 29.92 | 52.26 | 60.92 | 22.0 | 31.65 | 41.3 |
| Picea abies | 390 | 105 | 299 | 40 | 51 | 0 | 7.51 | 36.16 | 83.17 | 5.4 | 22.91 | 40.6 |
| Picea sitchensis | 19 | 6 | 19 | 0 | 0 | 0 | 7.67 | 30.46 | 58.31 | 5.4 | 20.16 | 29.9 |
| Pinus halepensis | 159 | 53 | 50 | 44 | 65 | 0 | 7.48 | 30.20 | 69.42 | 3.7 | 12.49 | 21.6 |
| Pinus laricio | 170 | 49 | 113 | 14 | 43 | 0 | 7.96 | 34.73 | 90.65 | 3.2 | 18.48 | 36.2 |
| Pinus nigra nigra | 114 | 34 | 43 | 22 | 49 | 0 | 7.64 | 29.39 | 63.38 | 5.4 | 16.13 | 25.7 |
| Pinus pinaster | 718 | 225 | 466 | 30 | 222 | 0 | 7.48 | 32.53 | 81.42 | 3.9 | 17.15 | 35.6 |
| Pinus pinea | 6 | 3 | 6 | 0 | 0 | 0 | 21.96 | 32.89 | 40.23 | 4.8 | 7.40 | 9.8 |
| Pinus sylvestris | 496 | 155 | 220 | 80 | 196 | 0 | 7.51 | 28.97 | 72.73 | 3.1 | 14.49 | 32.2 |
| Pinus uncinata | 25 | 10 | 9 | 9 | 7 | 0 | 7.93 | 26.15 | 56.18 | 2.5 | 9.66 | 20.2 |
| Populus alba | 11 | 1 | 0 | 0 | 11 | 0 | 26.93 | 73.82 | 92.79 | 13.5 | 31.75 | 39.0 |
| Populus nigra | 14 | 6 | 0 | 13 | 1 | 0 | 7.51 | 49.28 | 133.05 | 4.4 | 21.00 | 32.9 |
| Populus tremula | 29 | 11 | 4 | 9 | 16 | 0 | 8.59 | 26.20 | 83.33 | 8.1 | 21.21 | 31.9 |
| Pseudotsuga menziesii | 475 | 121 | 295 | 12 | 168 | 0 | 7.58 | 40.71 | 87.31 | 5.2 | 27.58 | 49.5 |
| Quercus cerris | 8 | 2 | 0 | 0 | 8 | 0 | 9.87 | 37.50 | 67.86 | 13.9 | 21.44 | 29.3 |
| Quercus ilex | 279 | 104 | 36 | 50 | 38 | 155 | 7.61 | 18.70 | 82.95 | 2.8 | 7.77 | 20.0 |
| Quercus petraea | 1,187 | 325 | 355 | 21 | 803 | 8 | 7.51 | 40.18 | 128.44 | 3.6 | 22.32 | 45.0 |
| Quercus pubescens | 796 | 253 | 181 | 71 | 442 | 102 | 7.54 | 23.59 | 87.22 | 2.7 | 11.97 | 29.6 |
| Quercus pyrenaica | 24 | 10 | 13 | 1 | 8 | 2 | 10.09 | 28.56 | 74.61 | 6.4 | 13.00 | 19.8 |
| Quercus robur | 1,121 | 323 | 235 | 64 | 817 | 5 | 7.48 | 42.19 | 133.69 | 3.6 | 20.66 | 40.3 |
| Quercus rubra | 49 | 13 | 14 | 0 | 35 | 0 | 7.58 | 28.98 | 88.11 | 9.0 | 20.98 | 34.6 |
| Quercus suber | 53 | 17 | 5 | 15 | 26 | 7 | 8.75 | 29.84 | 86.87 | 3.0 | 8.24 | 15.7 |
| Robinia pseudoacacia | 107 | 39 | 41 | 4 | 59 | 3 | 7.80 | 21.36 | 63.57 | 4.8 | 17.31 | 30.2 |
| Salix alba | 4 | 2 | 0 | 1 | 3 | 0 | 10.03 | 19.72 | 38.93 | 12.0 | 14.07 | 16.4 |
| Tilia cordata | 6 | 3 | 1 | 0 | 5 | 0 | 11.27 | 32.58 | 56.66 | 8.3 | 21.03 | 27.9 |
| Tilia platyphyllos | 4 | 2 | 2 | 0 | 2 | 0 | 8.47 | 19.31 | 31.86 | 8.8 | 12.72 | 16.4 |
| Ulmus minor | 13 | 6 | 2 | 3 | 8 | 0 | 8.09 | 13.72 | 32.91 | 3.7 | 8.25 | 15.5 |

### C. Practical example for the local recalibration method

We provide here a practical example of local recalibration for a given stand. We based this example on the Fagus sylvatica pure stand of figure 1, and partially detailed in table S4 below.

To perform a local recalibration of our height-diameter model, we selected four trees: three with the largest diameters (trees IDs 1 to 3) and one with the smallest diameter (tree ID 4). We computed predicted heights using only fixed effects, based on equation 4 (from main text) and Fagus sylvatica model parameter (table S7), for the 4 trees selected. The site-specific random effect was set to 0. This gave the simplified equation (4'):

$$\widehat{h_{0,ij}} = 1{,}3 + a_{s1} * (1 + a_{s,UA} + a_{s,CWS} + a_{s,C}) * (1 + a_{s2} * BA) * (1 - \exp(-a_{s3} * D_g{}^{a_{s4}})) * \frac{1}{1 + \frac{b_{s1} * (1 + b_{s,UA} + b_{s,CWS} + b_{s,C}) * (\exp(-b_{s2} * BA))}{\left(\frac{dbh}{D_g}\right)^{c_s}}} \quad (4')$$

*Table S 4: Fixed effects only predicted height*

| Tree_id | dbh (cm) | BA (cm) | Dg (cm) | Structure | Measured height ($h_{tot,ij}$) | Fixed effect prediction ($\widehat{h_{0,ij}}$) |
|---|---|---|---|---|---|---|
| 1 | 38.52 | 25.45 | 22.03 | Even-aged | 25 | 22.32 |
| 2 | 31.51 | 25.45 | 22.03 | Even-aged | 24.5 | 21.25 |
| 3 | 29.60 | 25.45 | 22.03 | Even-aged | 24 | 20.87 |
| 4 | 12.1 | 25.45 | 22.03 | Even-aged | 16.5 | 13.39 |

Using this data, we estimated the local site parameter $\lambda_{m,j}$ for the plot, using equation 6:

$$\lambda_{m,j} = \frac{1}{N_j} \sum \left( \frac{h_{tot,ij} - 1{,}3}{\widehat{h_{0,ij}} - 1{,}3} \right) - 1 \quad (6)$$

For this plot, $\lambda_{m,j} \approx 0.177$

Incorporating this local correction into equation 4 of the manuscript, we can calculate predicted heights after local recalibration. As a result, the root-mean square error, for the remaining 38 trees, decreases from 3.70m (fixed effects only) to 1.76m with local recalibration based on 4 trees.

### D. Model illustration for the 10 main species

*Figure S 2: Abies alba model, for varying basal area, quadratic diameter and stand characteristics. Predictions are limited to the range of diameter values observed for each structure in the calibration dataset.*

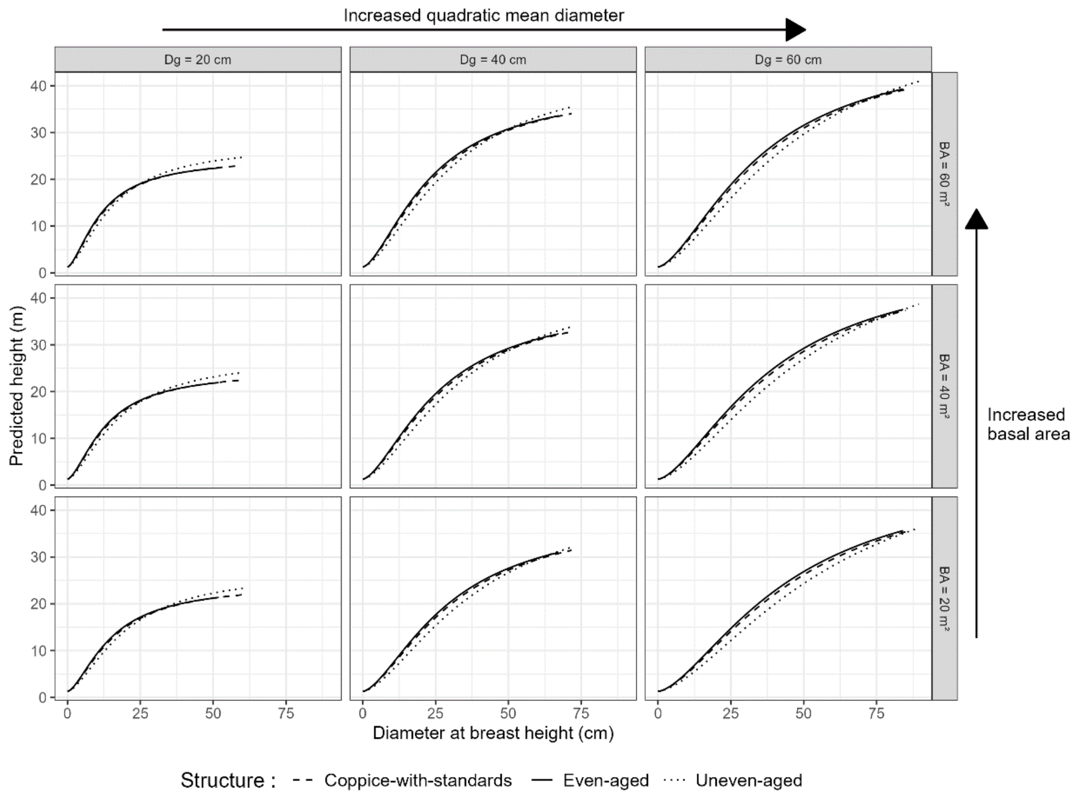


*Figure S 3: Castanea sativa model, for varying basal area, quadratic diameter and stand characteristics. Predictions are limited to the range of diameter values observed for each structure in the calibration dataset.*

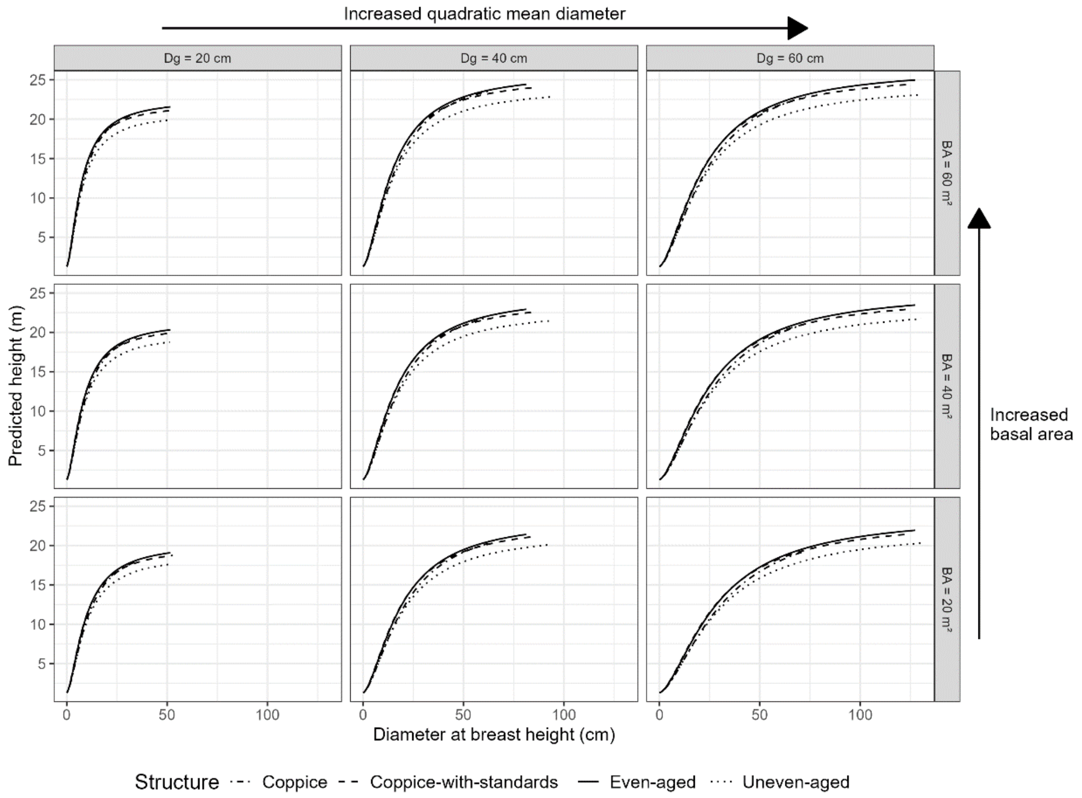

*Figure S 4: Fagus sylvatica model, for varying basal area, quadratic diameter and stand characteristics. Predictions are limited to the range of diameter values observed for each structure in the calibration dataset.*

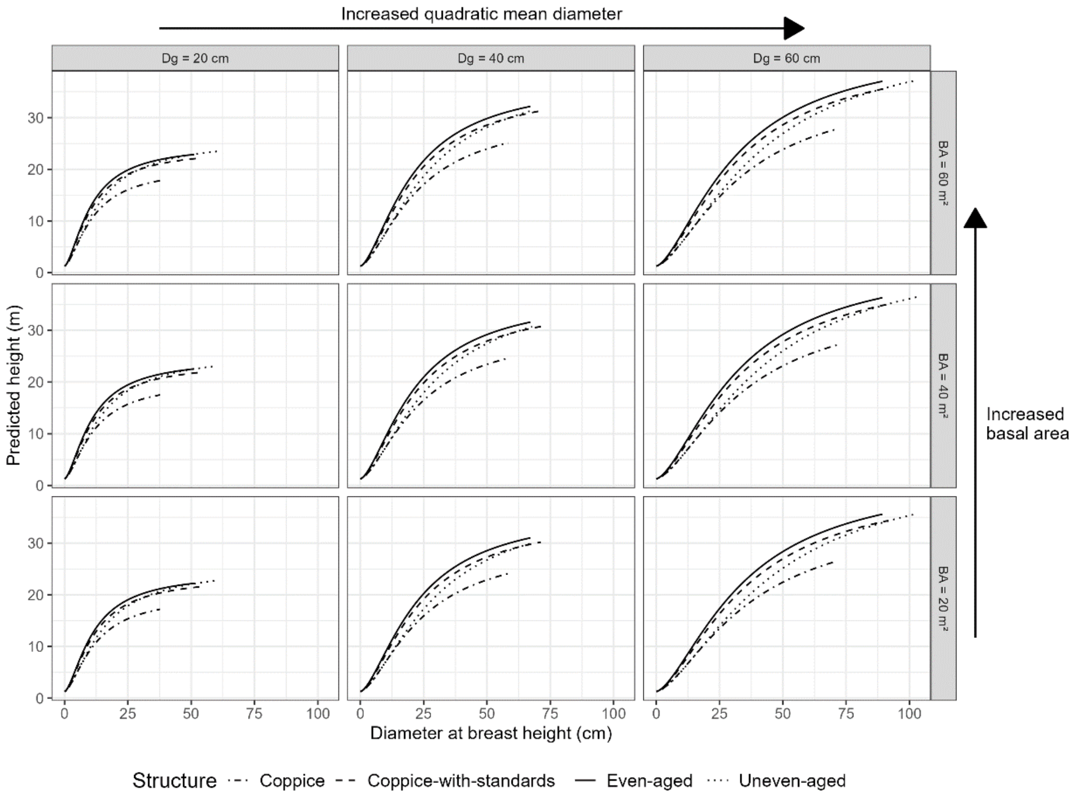


*Figure S 5: Picea abies model, for varying basal area, quadratic diameter and stand characteristics. Predictions are limited to the range of diameter values observed for each structure in the calibration dataset.*

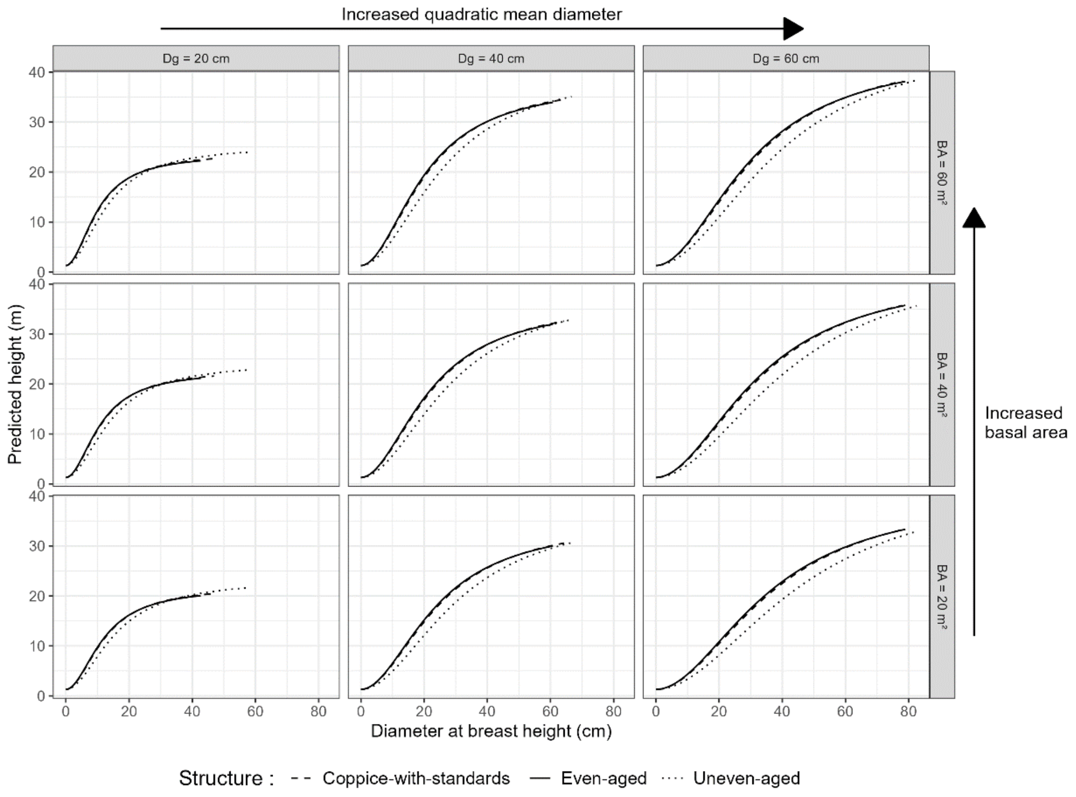

*Figure S 6: Pinus pinaster model, for varying basal area, quadratic diameter and stand characteristics. Predictions are limited to the range of diameter values observed for each structure in the calibration dataset.*

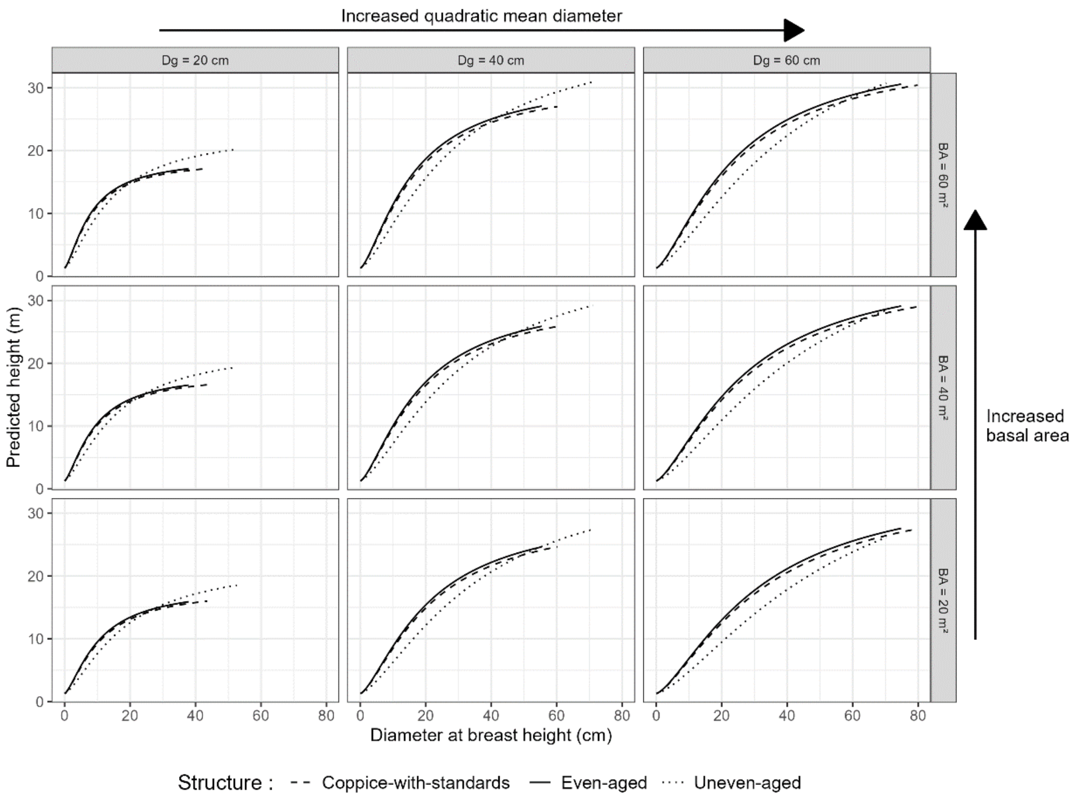


*Figure S 7: Pinus sylvestris model, for varying basal area, quadratic diameter and stand characteristics. Predictions are limited to the range of diameter values observed for each structure in the calibration dataset.*

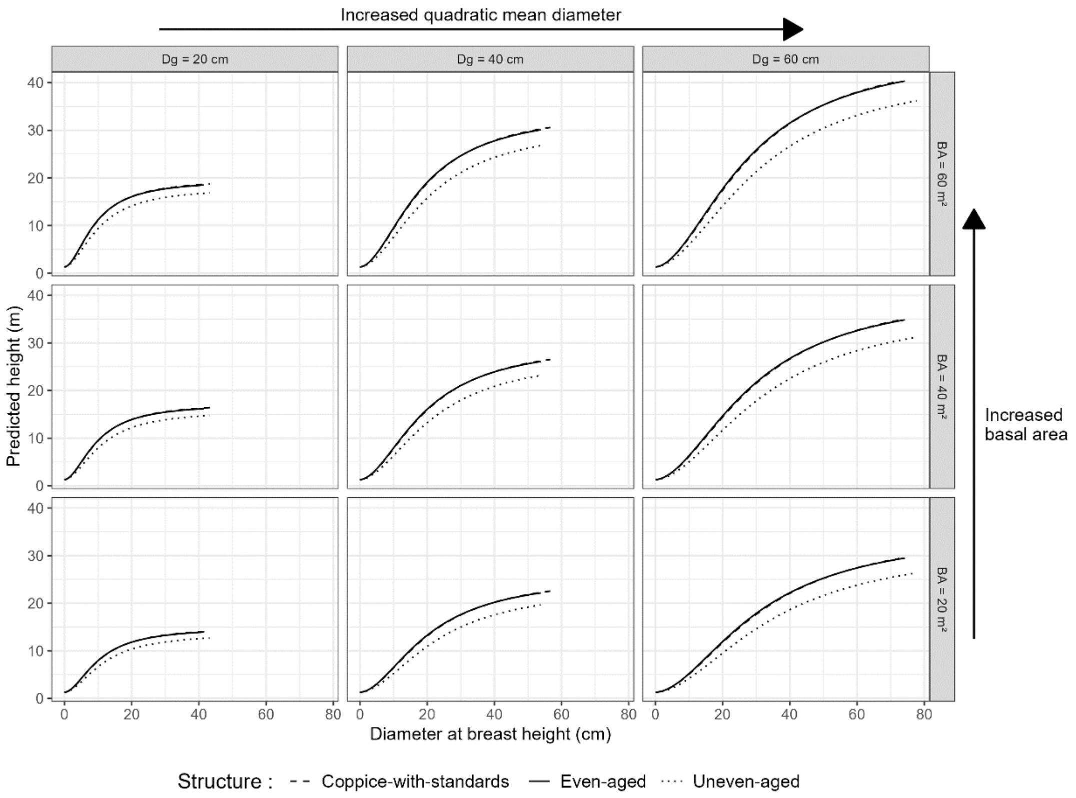

*Figure S 8: Pseudotsuga menziesii model, for varying basal area, quadratic diameter and stand characteristics. Predictions are limited to the range of diameter values observed for each structure in the calibration dataset.*

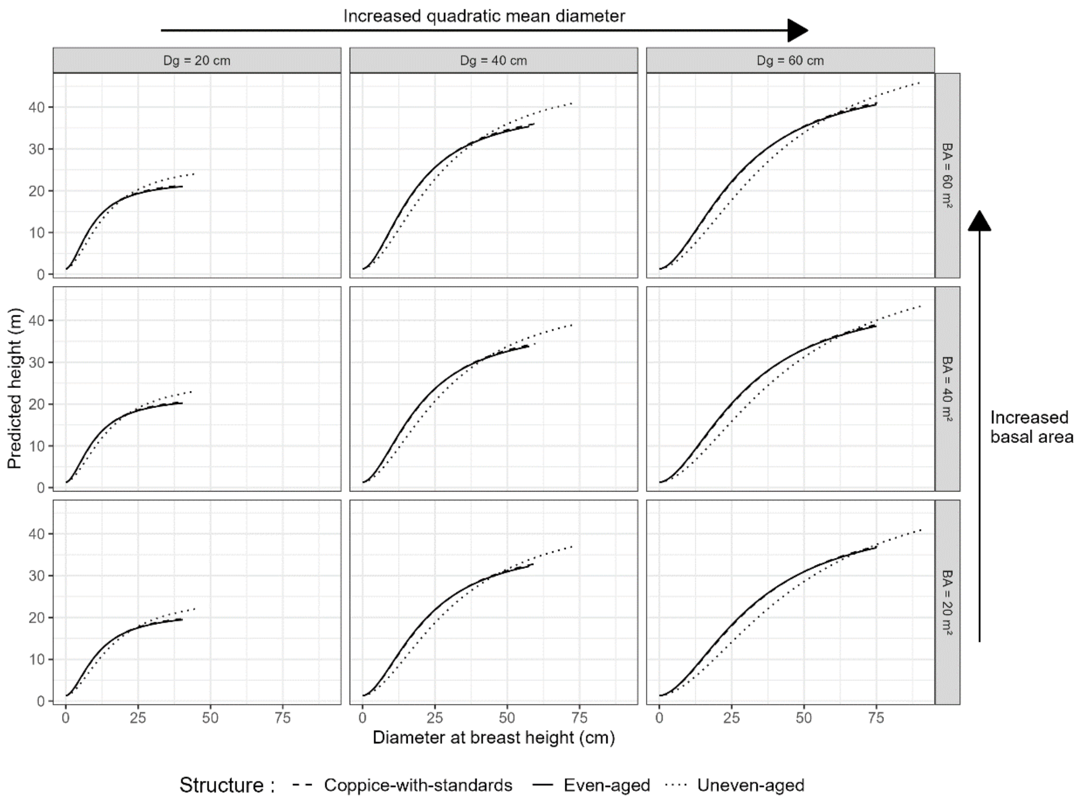


*Figure S 9: Quercus petraea model, for varying basal area, quadratic diameter and stand characteristics. Predictions are limited to the range of diameter values observed for each structure in the calibration dataset.*

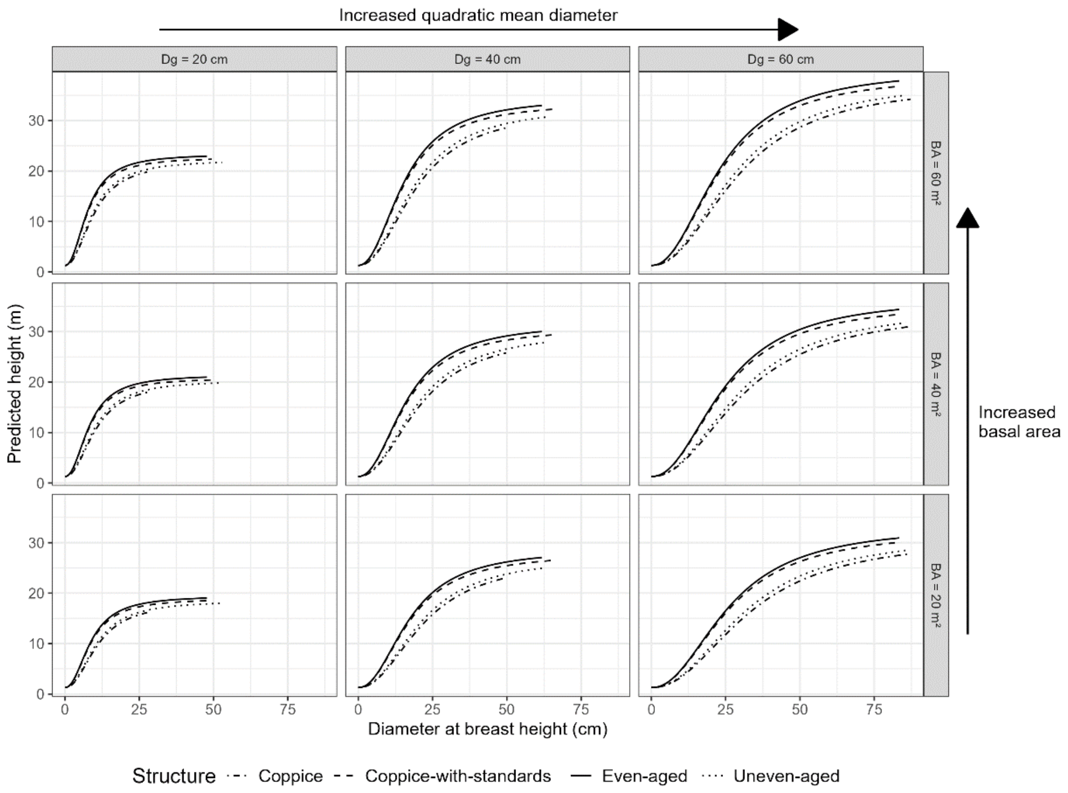

*Figure S 10: Quercus pubscens model, for varying basal area, quadratic diameter and stand characteristics. Predictions are limited to the range of diameter values observed for each structure in the calibration dataset.*

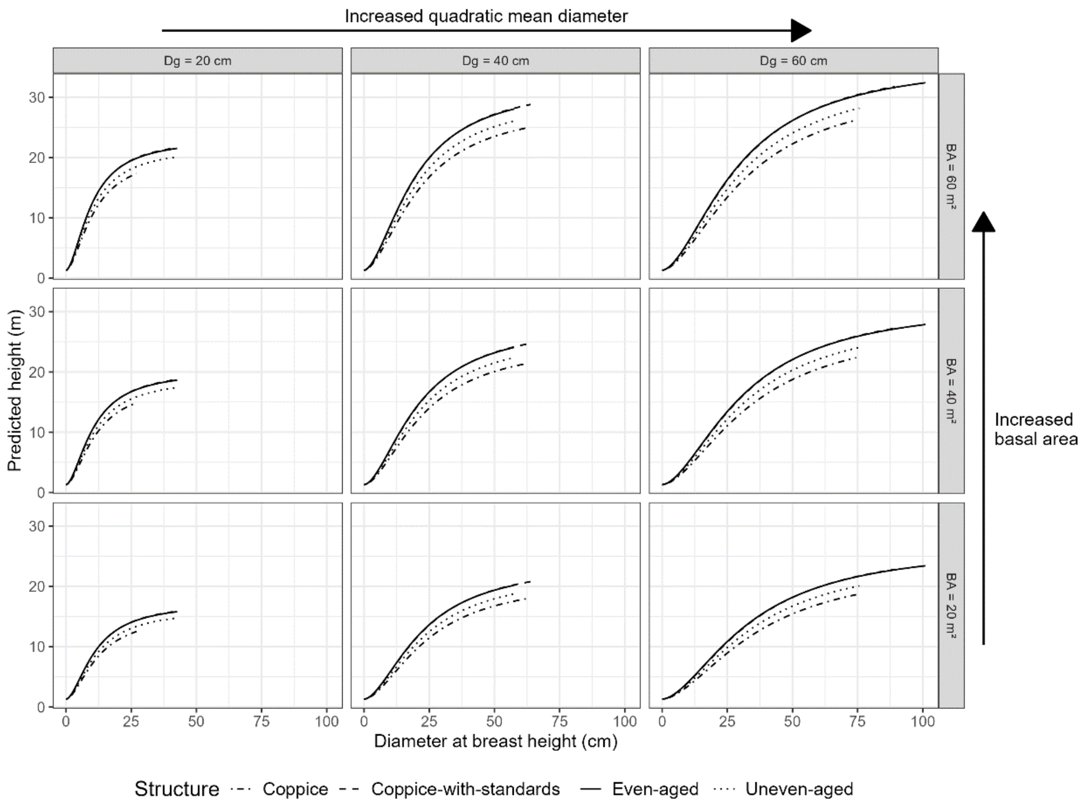


*Figure S 11: Quercus robur model, for varying basal area, quadratic diameter and stand characteristics. Predictions are limited to the range of diameter values observed for each structure in the calibration dataset.*

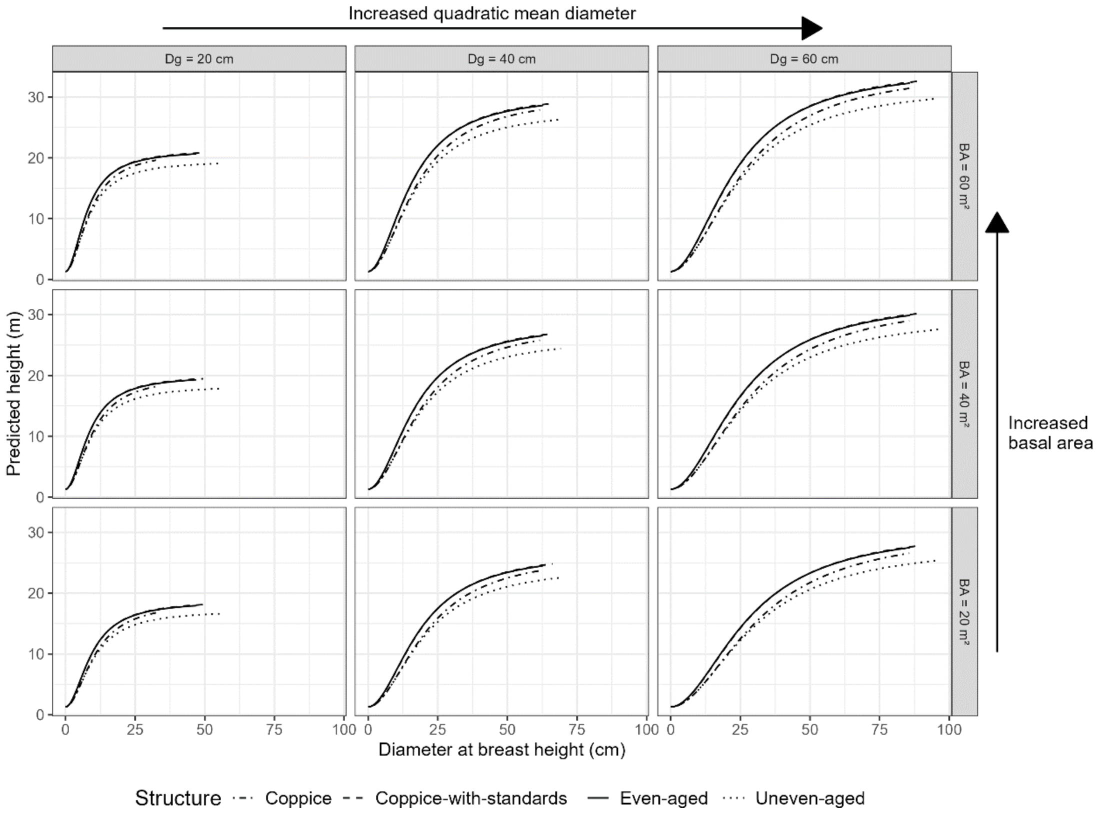

### E. Parameters of the models for all species

*Table S 5: Results of model calibration*

| Species | Estimation and Statistics | | | | | | | | | | | | | | Residual deviation | Deviation of random effects |
|---|---|---|---|---|---|---|---|---|---|---|---|---|---|---|---|---|
| | parameter | $a_{s1}$ | $a_{s,UA}$ | $a_{s,CWS}$ | $a_{s,C}$ | $a_{s2}$ | $a_{s3}$ | $a_{s4}$ | $b_{s1}$ | $b_{s,UA}$ | $b_{s,CWS}$ | $b_{s,C}$ | $b_{s2}$ | $c_s$ | | |
| Abies alba | estimate | 59.27 | 0.1295 | 0.0106 | - | 0.0003 | 0.0200 | 1.0637 | 0.7621 | 0.5553 | 0.0832 | - | 0.0094 | 1.5730 | 2.04 | 0.13 |
| | significativity | *** | *** | . | - | . | *** | *** | *** | *** | *** | - | *** | *** | | |
| | stddev | 2.64 | 0.01113 | 0.01003 | - | 0.00027 | 0.00108 | 0.03178 | 0.02310 | 0.02907 | 0.02282 | - | 0.00051 | 0.01615 | | |
| Abies grandis | estimate | 177.87 | - | 0.01484 | - | -0.00059 | 0.00777 | 1.04155 | 1.82638 | - | -0.12775 | - | 0.01510 | 0.86069 | 1.59 | 0.13 |
| | significativity | . | - | . | - | . | . | *** | . | - | . | - | * | *** | | |
| | stddev | 310.98 | - | 0.1563 | - | 0.0040 | 0.0090 | 0.2423 | 1.2016 | - | 0.2286 | - | 0.0063 | 0.1998 | | |
| Acer campestre | estimate | 29.74 | 0.1788 | -0.0043 | 0.0038 | 0.0112 | 0.0622 | 0.7509 | 0.3877 | 0.4593 | -0.1464 | -0.3519 | 0.0072 | 1.8099 | 2.12 | 0.19 |
| | significativity | . | . | . | . | *** | * | ** | *** | . | . | . | . | *** | | |
| | stddev | 24.25 | 0.0988 | 0.0598 | 0.1177 | 0.0033 | 0.0266 | 0.2675 | 0.1117 | 0.3099 | 0.1421 | 0.3107 | 0.0066 | 0.2403 | | |
| Acer pseudoplatanus | estimate | 39.57 | 0.0218 | 0.0469 | -0.1736 | 0.0050 | 0.0431 | 0.9289 | 0.5078 | 0.1282 | 0.0668 | -0.3040 | 0.0129 | 1.4940 | 2.74 | 0.16 |
| | significativity | *** | . | . | * | * | *** | *** | *** | . | . | . | ** | *** | | |
| | stddev | 7.86 | 0.1116 | 0.0409 | 0.0725 | 0.0020 | 0.0066 | 0.1251 | 0.0996 | 0.3374 | 0.1040 | 0.1863 | 0.0041 | 0.1330 | | |
| Alnus glutinosa | estimate | 30.39 | -0.0754 | -0.0127 | -0.0618 | 0.0034 | 0.0896 | 0.8291 | 0.2963 | 0.2875 | 0.2225 | 0.1303 | 0.0022 | 1.7421 | 2.27 | 0.17 |
| | significativity | *** | . | . | . | *** | *** | *** | *** | . | ** | . | . | *** | | |
| | stddev | 3.25 | 0.0410 | 0.0242 | 0.0337 | 0.0008 | 0.0154 | 0.1128 | 0.0343 | 0.1705 | 0.0766 | 0.1071 | 0.0015 | 0.0834 | | |
| Betula pendula | estimate | 23.16 | -0.0264 | -0.0059 | -0.0673 | 0.0149 | 0.1014 | 0.8368 | 0.2721 | 0.1619 | 0.0384 | -0.0304 | -0.0043 | 1.8680 | 1.56 | 0.16 |
| | significativity | *** | . | . | ** | *** | *** | *** | *** | . | . | . | * | *** | | |
| | stddev | 2.13 | 0.0420 | 0.0207 | 0.0254 | 0.0015 | 0.0147 | 0.1061 | 0.0271 | 0.1549 | 0.0589 | 0.0732 | 0.0020 | 0.0911 | | |
| Betula pubescens | estimate | 77.2 | -0.1847 | 0.0020 | -0.0500 | -0.0010 | 0.0734 | 0.4998 | 0.5799 | -0.2212 | 0.2312 | 0.7933 | 0.0416 | 2.0110 | 1.17 | 0.17 |
| | significativity | . | ** | . | . | . | . | . | *** | . | . | * | *** | *** | | |
| | stddev | 571.21 | 0.0692 | 0.0676 | 0.0984 | 0.0033 | 0.4910 | 0.7046 | 0.1594 | 0.1862 | 0.1975 | 0.3939 | 0.0085 | 0.2463 | | |
| Carpinus betulus | estimate | 33.32 | -0.1015 | -0.0028 | -0.0814 | 0.0069 | 0.0622 | 0.9149 | 0.3978 | -0.0923 | 0.0732 | -0.0744 | 0.0015 | 1.5109 | 1.98 | 0.16 |
| | significativity | *** | ** | . | * | *** | *** | *** | *** | . | . | . | . | *** | | |
| | stddev | 2.79 | 0.0392 | 0.0159 | 0.0358 | 0.0009 | 0.0060 | 0.0726 | 0.0352 | 0.1324 | 0.0463 | 0.1056 | 0.0019 | 0.0619 | | |

| Species | Estimation and Statistics | | | | | | | | | | | | | | Residual deviation | Deviation of random effects |
|---|---|---|---|---|---|---|---|---|---|---|---|---|---|---|---|---|
| | parameter | $a_{s1}$ | $a_{s,UA}$ | $a_{s,CWS}$ | $a_{s,C}$ | $a_{s2}$ | $a_{s3}$ | $a_{s4}$ | $b_{s1}$ | $b_{s,UA}$ | $b_{s,CWS}$ | $b_{s,C}$ | $b_{s2}$ | $c_s$ | | |
| Castanea sativa | estimate | 21.36 | -0.0798 | -0.0260 | 0.0096 | 0.0031 | 0.0762 | 1.0608 | 0.3808 | 0.0305 | -0.0671 | 0.1576 | 0.0097 | 1.5726 | 1.67 | 0.21 |
| | significativity | *** | ** | * | . | *** | *** | *** | *** | . | * | *** | *** | *** | | |
| | stddev | 0.52 | 0.02736 | 0.01280 | 0.01654 | 0.00045 | 0.00926 | 0.05363 | 0.02140 | 0.08879 | 0.03017 | 0.04726 | 0.00085 | 0.03545 | | |
| Cedrus atlantica | estimate | 60.97 | -0.09766 | 0.04304 | - | -0.00366 | 0.01334 | 1.24083 | 2.05717 | -0.28479 | 0.04620 | - | 0.02261 | 1.06650 | 1.25 | 0.19 |
| | significativity | *** | . | . | - | . | *** | *** | ** | . | . | - | *** | *** | | |
| | stddev | 17.37 | 0.1856 | 0.1543 | - | 0.0019 | 0.0035 | 0.1363 | 0.7091 | 0.2316 | 0.2431 | - | 0.0034 | 0.1384 | | |
| Corylus avellana | estimate | 50.8 | -0.3915 | -0.3556 | -0.0293 | -0.0086 | 0.1604 | 0.9375 | 4.7312 | -0.5231 | -0.4938 | -0.0181 | 0.0263 | 0.3161 | 0.79 | 0.16 |
| | significativity | . | . | . | . | * | . | . | . | . | . | . | ** | . | | |
| | stddev | 104.23 | 0.7880 | 0.7935 | 1.0665 | 0.0041 | 0.2383 | 1.1460 | 11.3395 | 0.7321 | 0.7394 | 1.3744 | 0.0089 | 0.1948 | | |
| Fagus sylvatica | estimate | 49.75 | 0.0412 | -0.0338 | -0.1867 | 0.0005 | 0.0318 | 0.9863 | 0.4410 | 0.5103 | 0.0530 | 0.1154 | 0.0029 | 1.5511 | 2.38 | 0.19 |
| | significativity | *** | * | *** | *** | . | *** | *** | *** | *** | ** | . | *** | *** | | |
| | stddev | 1.48 | 0.01789 | 0.00757 | 0.01948 | 0.00028 | 0.00134 | 0.02421 | 0.01293 | 0.05093 | 0.01709 | 0.06320 | 0.00053 | 0.01841 | | |
| Fraxinus excelsior | estimate | 34.39 | -0.16167 | 0.00545 | -0.07635 | 0.00847 | 0.04185 | 1.02839 | 0.32658 | -0.24566 | 0.04709 | 0.03083 | -0.00613 | 1.65759 | 2.94 | 0.18 |
| | significativity | *** | *** | . | * | *** | *** | *** | *** | *** | . | . | *** | *** | | |
| | stddev | 1.81 | 0.02372 | 0.01611 | 0.03206 | 0.00084 | 0.00404 | 0.05705 | 0.02106 | 0.06062 | 0.03640 | 0.09099 | 0.00114 | 0.04454 | | |
| Larix decidua | estimate | 41.8 | -0.02892 | 0.03465 | - | 0.00224 | 0.02661 | 1.02720 | 0.59842 | 0.14805 | -0.02910 | - | 0.01074 | 1.80725 | 2.17 | 0.15 |
| | significativity | *** | . | . | - | ** | *** | *** | *** | * | . | - | *** | *** | | |
| | stddev | 4.65 | 0.02698 | 0.03719 | - | 0.00086 | 0.00362 | 0.08292 | 0.05191 | 0.06699 | 0.08500 | - | 0.00155 | 0.06739 | | |
| Larix kaempferi | estimate | 36.16 | - | -0.01465 | - | 0.01150 | 0.01419 | 1.17270 | 0.13076 | - | 0.15636 | - | -0.00051 | 3.23541 | 0.63 | 0.12 |
| | significativity | ** | - | . | - | *** | *** | *** | *** | - | . | - | . | *** | | |
| | stddev | 13.47 | - | 0.04979 | - | 0.00314 | 0.00353 | 0.22630 | 0.03680 | - | 0.23589 | - | 0.00532 | 0.45188 | | |
| Picea abies | estimate | 39.91 | 0.07737 | 0.01371 | - | 0.00246 | 0.01509 | 1.26384 | 0.43086 | 0.65289 | 0.06797 | - | 0.00797 | 1.93804 | 1.92 | 0.15 |
| | significativity | *** | *** | . | - | *** | *** | . | *** | *** | * | - | *** | *** | | |
| | stddev | 1.06 | 0.01273 | 0.01118 | - | 0.00028 | 0.00107 | 0.03250 | 0.01399 | 0.03865 | 0.02768 | - | 0.00049 | 0.02503 | | |
| Picea sitchensis | estimate | 153.87 | -0.57289 | -0.37316 | - | -0.00398 | 0.00610 | 1.52234 | 5.44565 | -0.69936 | -0.51670 | - | 0.01240 | 0.54782 | 1.96 | 0.15 |
| | significativity | . | ** | * | - | . | *** | *** | . | *** | *** | - | ** | *** | | |
| | stddev | 89.71 | 0.1945 | 0.1580 | - | 0.0026 | 0.0018 | 0.1218 | 3.8946 | 0.1904 | 0.1382 | - | 0.0041 | 0.0833 | | |

| Species | Estimation and Statistics | | | | | | | | | | | | | | Residual deviation | Deviation of random effects |
|---|---|---|---|---|---|---|---|---|---|---|---|---|---|---|---|---|
| | parameter | $a_{s1}$ | $a_{s,UA}$ | $a_{s,CWS}$ | $a_{s,C}$ | $a_{s2}$ | $a_{s3}$ | $a_{s4}$ | $b_{s1}$ | $b_{s,UA}$ | $b_{s,CWS}$ | $b_{s,C}$ | $b_{s2}$ | $c_s$ | | |
| Pinus halepensis | estimate | 61.46 | 0.1080 | 0.1288 | - | -0.0052 | 0.0398 | 0.7730 | 1.7510 | 0.2769 | 0.2109 | - | 0.0331 | 0.8301 | 1.84 | 0.16 |
| | significativity | ** | . | *** | - | *** | *** | *** | *** | * | ** | - | *** | *** | | |
| | stddev | 21.38 | 0.06182 | 0.03796 | - | 0.00132 | 0.00682 | 0.08580 | 0.25152 | 0.11554 | 0.07047 | - | 0.00232 | 0.04908 | | |
| Pinus laricio | estimate | 27.34 | 0.06162 | 0.03744 | - | 0.00639 | 0.00777 | 1.50161 | 0.26960 | 0.47896 | 0.10130 | - | -0.00300 | 1.62899 | 1.49 | 0.20 |
| | significativity | *** | . | . | - | *** | *** | *** | *** | *** | * | - | ** | *** | | |
| | stddev | 1.2 | 0.03986 | 0.02249 | - | 0.00085 | 0.00102 | 0.05383 | 0.01784 | 0.07964 | 0.05134 | - | 0.00091 | 0.05029 | | |
| Pinus nigra nigra | estimate | 29.64 | 0.08400 | 0.00590 | - | 0.00466 | 0.01207 | 1.32803 | 0.49353 | 0.44737 | 0.08065 | - | 0.00928 | 1.52525 | 2.29 | 0.18 |
| | significativity | *** | * | . | - | *** | *** | *** | *** | *** | . | - | *** | *** | | |
| | stddev | 2.28 | 0.03675 | 0.02123 | - | 0.00078 | 0.00147 | 0.06887 | 0.04290 | 0.10143 | 0.05518 | - | 0.00123 | 0.06091 | | |
| Pinus pinaster | estimate | 35.06 | 0.23787 | -0.01374 | - | 0.00092 | 0.00967 | 1.40117 | 0.47802 | 1.17399 | 0.04229 | - | 0.00921 | 1.42384 | 1.18 | 0.15 |
| | significativity | *** | *** | . | - | ** | *** | *** | *** | *** | . | - | *** | *** | | |
| | stddev | 0.71 | 0.02923 | 0.00899 | - | 0.00030 | 0.00036 | 0.01761 | 0.01922 | 0.08918 | 0.02406 | - | 0.00065 | 0.02941 | | |
| Pinus pinea | estimate | 72.01 | - | 0.55781 | - | -0.00797 | 0.00967 | 1.14314 | 2.81426 | - | 0.87597 | - | 0.03419 | 0.59926 | 0.86 | 0.16 |
| | significativity | . | - | . | - | . | *** | *** | . | - | . | - | ** | *** | | |
| | stddev | 51.86 | - | 0.4933 | - | 0.0069 | 0.0023 | 0.1552 | 2.0367 | - | 0.6630 | - | 0.0118 | 0.1464 | | |
| Pinus sylvestris | estimate | 103.55 | -0.08569 | 0.01118 | - | 0.01033 | 0.00743 | 0.91605 | 0.32923 | 0.26842 | 0.04402 | - | 0.00518 | 1.81832 | 2.32 | 0.25 |
| | significativity | . | *** | . | - | *** | . | *** | *** | *** | . | - | *** | *** | | |
| | stddev | 86.87 | 0.01670 | 0.01188 | - | 0.00063 | 0.00534 | 0.06061 | 0.01348 | 0.05030 | 0.02640 | - | 0.00072 | 0.03242 | | |
| Pinus uncinata | estimate | 15.16 | 0.08001 | 0.05561 | - | 0.00407 | 0.00825 | 1.70911 | 0.47089 | 0.49247 | -0.00913 | - | 0.00711 | 1.83750 | 1.74 | 0.20 |
| | significativity | *** | * | . | - | ** | * | *** | *** | *** | . | - | ** | *** | | |
| | stddev | 1.05 | 0.04037 | 0.05235 | - | 0.00155 | 0.00367 | 0.18032 | 0.06083 | 0.10326 | 0.11008 | - | 0.00218 | 0.09260 | | |
| Populus alba | estimate | 54.02 | - | -0.22960 | - | -0.00159 | 0.02986 | 1.01660 | 0.67232 | - | -0.53816 | - | 0.00620 | 1.65439 | 0.84 | 0.13 |
| | significativity | * | - | . | - | . | . | ** | * | - | *** | - | . | *** | | |
| | stddev | 20.97 | - | 0.1207 | - | 0.0017 | 0.0232 | 0.3765 | 0.2911 | - | 0.1208 | - | 0.0043 | 0.2915 | | |
| Populus nigra | estimate | 45.55 | 0.01908 | 0.17718 | 0.00533 | 0.00146 | 0.04281 | 0.89453 | 0.55037 | 0.22558 | 0.32823 | 0.13961 | 0.00649 | 1.50156 | 1.67 | 0.20 |
| | significativity | *** | . | . | . | . | *** | *** | ** | . | . | . | . | *** | | |
| | stddev | 9.25 | 0.1062 | 0.1013 | 0.1622 | 0.0027 | 0.0121 | 0.1288 | 0.1763 | 0.2647 | 0.2251 | 0.4068 | 0.0052 | 0.2092 | | |

| Species | Estimation and Statistics parameter | $a_{s1}$ | $a_{s,UA}$ | $a_{s,CWS}$ | $a_{s,C}$ | $a_{s2}$ | $a_{s3}$ | $a_{s4}$ | $b_{s1}$ | $b_{s,UA}$ | $b_{s,CWS}$ | $b_{s,C}$ | $b_{s2}$ | $c_s$ | Residual deviation | Deviation of random effects |
|---|---|---|---|---|---|---|---|---|---|---|---|---|---|---|---|---|
| Populus tremula | estimate | 37.46 | -0.0570 | 0.0059 | -0.0455 | 0.0034 | 0.0528 | 1.0280 | 0.6172 | 0.0139 | 0.0779 | -0.0333 | 0.0060 | 1.3176 | | |
| | significativity | *** | . | . | . | . | *** | *** | *** | . | . | . | . | *** | 1.45 | 0.16 |
| | stddev | 4.84 | 0.07281 | 0.05081 | 0.06852 | 0.00202 | 0.01109 | 0.11719 | 0.14178 | 0.19767 | 0.12698 | 0.16188 | 0.00401 | 0.12978 | | |
| Pseudotsuga menziesii | estimate | 45.07 | 0.20845 | 0.01852 | - | 0.00160 | 0.00718 | 1.45561 | 0.39143 | 0.97330 | 0.05445 | - | 0.00652 | 1.74230 | | |
| | significativity | *** | *** | . | - | *** | *** | *** | *** | *** | . | - | *** | *** | 1.85 | 0.13 |
| | stddev | 1.01 | 0.0377 | 0.0111 | - | 0.0003 | 0.0004 | 0.0234 | 0.0153 | 0.1216 | 0.0299 | - | 0.0005 | 0.0338 | | |
| Quercus cerris | estimate | 24.7 | - | 1.5352 | -0.1650 | 0.0875 | 0.0165 | 1.3212 | 0.9010 | - | 2.7847 | -0.2254 | -0.0250 | 0.7409 | | |
| | significativity | ** | - | . | . | *** | . | *** | . | - | . | . | *** | *** | 3.07 | 0.09 |
| | stddev | 8.16 | - | 1.8458 | 0.2703 | 0.0234 | 0.0086 | 0.2138 | 0.5354 | - | 3.0787 | 0.4066 | 0.0027 | 0.0831 | | |
| Quercus ilex | estimate | 19.49 | 0.08149 | 0.02243 | -0.03173 | 0.00859 | 0.03850 | 1.09240 | 0.75498 | 0.57219 | 0.17995 | 0.46986 | 0.00785 | 1.01378 | | |
| | significativity | *** | . | . | . | *** | *** | *** | *** | * | . | *** | *** | *** | 1.07 | 0.22 |
| | stddev | 2.12 | 0.0856 | 0.0428 | 0.0454 | 0.0016 | 0.0044 | 0.0728 | 0.1195 | 0.2268 | 0.0981 | 0.1335 | 0.0019 | 0.0662 | | |
| Quercus petraea | estimate | 32.63 | -0.0534 | -0.0294 | -0.0721 | 0.0060 | 0.0330 | 1.0137 | 0.1874 | 0.5170 | -0.0003 | 0.6569 | 0.0061 | 2.1660 | | |
| | significativity | *** | ** | *** | *** | *** | *** | *** | *** | *** | . | *** | *** | *** | 2.02 | 0.15 |
| | stddev | 0.54 | 0.01642 | 0.00470 | 0.01659 | 0.00032 | 0.00121 | 0.01770 | 0.00609 | 0.08860 | 0.01602 | 0.09265 | 0.00080 | 0.02605 | | |
| Quercus pubescens | estimate | 22.58 | -0.06542 | 0.00810 | -0.11512 | 0.01122 | 0.04416 | 0.99286 | 0.42095 | 0.06926 | 0.02891 | 0.16874 | 0.00583 | 1.70921 | | |
| | significativity | *** | ** | . | *** | *** | *** | *** | *** | . | . | *** | *** | *** | 1.27 | 0.24 |
| | stddev | 1.2 | 0.02237 | 0.01155 | 0.01390 | 0.00071 | 0.00293 | 0.04387 | 0.01796 | 0.06035 | 0.02370 | 0.04067 | 0.00090 | 0.03229 | | |
| Quercus pyrenaica | estimate | 21.97 | 0.02127 | 0.04240 | -0.00975 | 0.00940 | 0.02036 | 1.34827 | 0.46902 | 0.07001 | -0.06052 | -0.50964 | 0.00679 | 1.90208 | | |
| | significativity | *** | . | . | . | ** | ** | *** | *** | . | . | ** | . | *** | 1.97 | 0.15 |
| | stddev | 2.08 | 0.08925 | 0.04656 | 0.10558 | 0.00300 | 0.00693 | 0.14724 | 0.10249 | 0.25272 | 0.10165 | 0.19182 | 0.00564 | 0.17520 | | |
| Quercus robur | estimate | 30.27 | -0.08605 | 0.00685 | -0.00597 | 0.00396 | 0.03890 | 0.99302 | 0.27664 | 0.16480 | 0.02373 | 0.28099 | 0.00844 | 1.91118 | | |
| | significativity | *** | *** | . | . | *** | *** | *** | *** | ** | . | *** | *** | *** | 2.11 | 0.17 |
| | stddev | 0.55 | 0.01471 | 0.00620 | 0.02071 | 0.00033 | 0.00176 | 0.02094 | 0.00985 | 0.06260 | 0.01900 | 0.08086 | 0.00080 | 0.02957 | | |
| Quercus rubra | estimate | 28.58 | 0.03804 | 0.08733 | - | 0.01769 | 0.05212 | 0.94945 | 0.27130 | -0.07625 | 0.13770 | - | -0.01209 | 1.47452 | | |
| | significativity | *** | . | * | - | *** | *** | *** | *** | . | . | - | *** | *** | 1.42 | 0.16 |
| | stddev | 3.7 | 0.0860 | 0.0427 | - | 0.0030 | 0.0115 | 0.1303 | 0.0412 | 0.2009 | 0.0976 | - | 0.0032 | 0.1065 | | |

| Species | Estimation and Statistics parameter | $a_{s1}$ | $a_{s,UA}$ | $a_{s,CWS}$ | $a_{s,C}$ | $a_{s2}$ | $a_{s3}$ | $a_{s4}$ | $b_{s1}$ | $b_{s,UA}$ | $b_{s,CWS}$ | $b_{s,C}$ | $b_{s2}$ | $c_s$ | Residual deviation | Deviation of random effects |
|---|---|---|---|---|---|---|---|---|---|---|---|---|---|---|---|---|
| Quercus suber | estimate | 56.62 | 0.3080 | -0.1981 | -0.1834 | -0.0001 | 0.0211 | 0.9308 | 2.2459 | 0.4097 | -0.3096 | -0.0763 | 0.0035 | 0.8299 | 1.03 | 0.18 |
| | significativity | . | . | . | . | . | *** | *** | * | . | * | . | . | *** | | |
| | stddev | 29.65 | 0.3351 | 0.1107 | 0.1790 | 0.0043 | 0.0040 | 0.1397 | 0.9636 | 0.4794 | 0.1290 | 0.3035 | 0.0066 | 0.1080 | | |
| Robinia pseudoacacia | estimate | 26.61 | -0.0780 | 0.0309 | 0.0437 | 0.0055 | 0.0516 | 1.1228 | 0.3479 | -0.1100 | -0.0675 | 0.0568 | -0.0062 | 1.9347 | 2.08 | 0.17 |
| | significativity | *** | . | . | . | *** | *** | *** | *** | . | . | . | *** | *** | | |
| | stddev | 1.74 | 0.0440 | 0.0244 | 0.0318 | 0.0011 | 0.0100 | 0.1030 | 0.0323 | 0.1135 | 0.0501 | 0.0728 | 0.0016 | 0.0710 | | |
| Salix alba | estimate | 26.62 | -0.1326 | 0.1050 | -0.1048 | -0.0002 | 0.0493 | 1.1260 | 0.4430 | -0.0296 | 0.4050 | -0.1480 | 0.0096 | 1.6306 | 3.74 | 0.13 |
| | significativity | *** | . | . | . | . | * | *** | *** | . | . | . | * | *** | | |
| | stddev | 2.86 | 0.07872 | 0.08392 | 0.08707 | 0.00165 | 0.02297 | 0.18515 | 0.12611 | 0.26221 | 0.29607 | 0.28071 | 0.00462 | 0.19617 | | |
| Tilia cordata | estimate | 54.12 | - | 0.00774 | 0.08332 | -0.00883 | 0.01860 | 1.44491 | 2.27422 | - | 0.15941 | 0.21465 | 0.04036 | 0.90718 | 1.70 | 0.14 |
| | significativity | ** | - | . | . | * | . | *** | * | - | . | . | *** | *** | | |
| | stddev | 16.38 | - | 0.1274 | 0.2421 | 0.0037 | 0.0152 | 0.3315 | 1.1301 | - | 0.2608 | 0.5594 | 0.0099 | 0.1578 | | |
| Tilia platyphyllos | estimate | 18.77 | -0.05710 | -0.07229 | -0.13002 | 0.01098 | 0.03181 | 1.30644 | 0.13396 | 0.21984 | 0.25089 | 0.34147 | -0.01418 | 2.19324 | 2.57 | 0.17 |
| | significativity | *** | . | . | . | ** | . | * | ** | . | . | . | ** | *** | | |
| | stddev | 2.39 | 0.0995 | 0.0877 | 0.1353 | 0.0039 | 0.0459 | 0.5594 | 0.0439 | 0.2142 | 0.2223 | 0.6740 | 0.0046 | 0.2526 | | |
| Ulmus minor | estimate | 26.56 | -0.4122 | -0.1693 | -0.3720 | 0.0069 | 0.0371 | 1.2522 | 1.1906 | -0.6596 | -0.4163 | -0.6637 | 0.0176 | 1.7364 | 1.02 | 0.20 |
| | significativity | ** | *** | . | *** | . | . | ** | * | *** | ** | *** | * | *** | | |
| | stddev | 8.98 | 0.0984 | 0.1097 | 0.0958 | 0.0052 | 0.0254 | 0.4073 | 0.4992 | 0.1258 | 0.1278 | 0.0838 | 0.0082 | 0.3876 | | |

### F. Illustration of basal area effect

*Figure S 12: Correlation effect between basal area parameters. example of Tilia cordata species.*

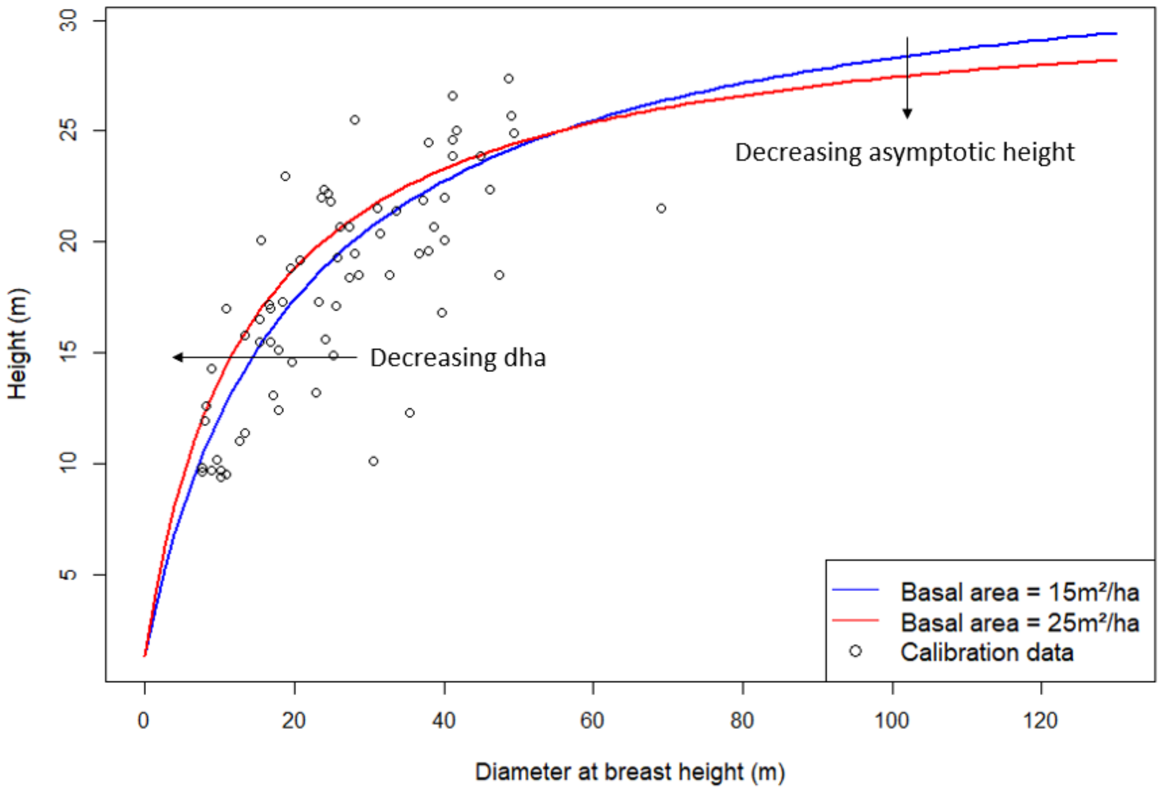

### G. Model residuals

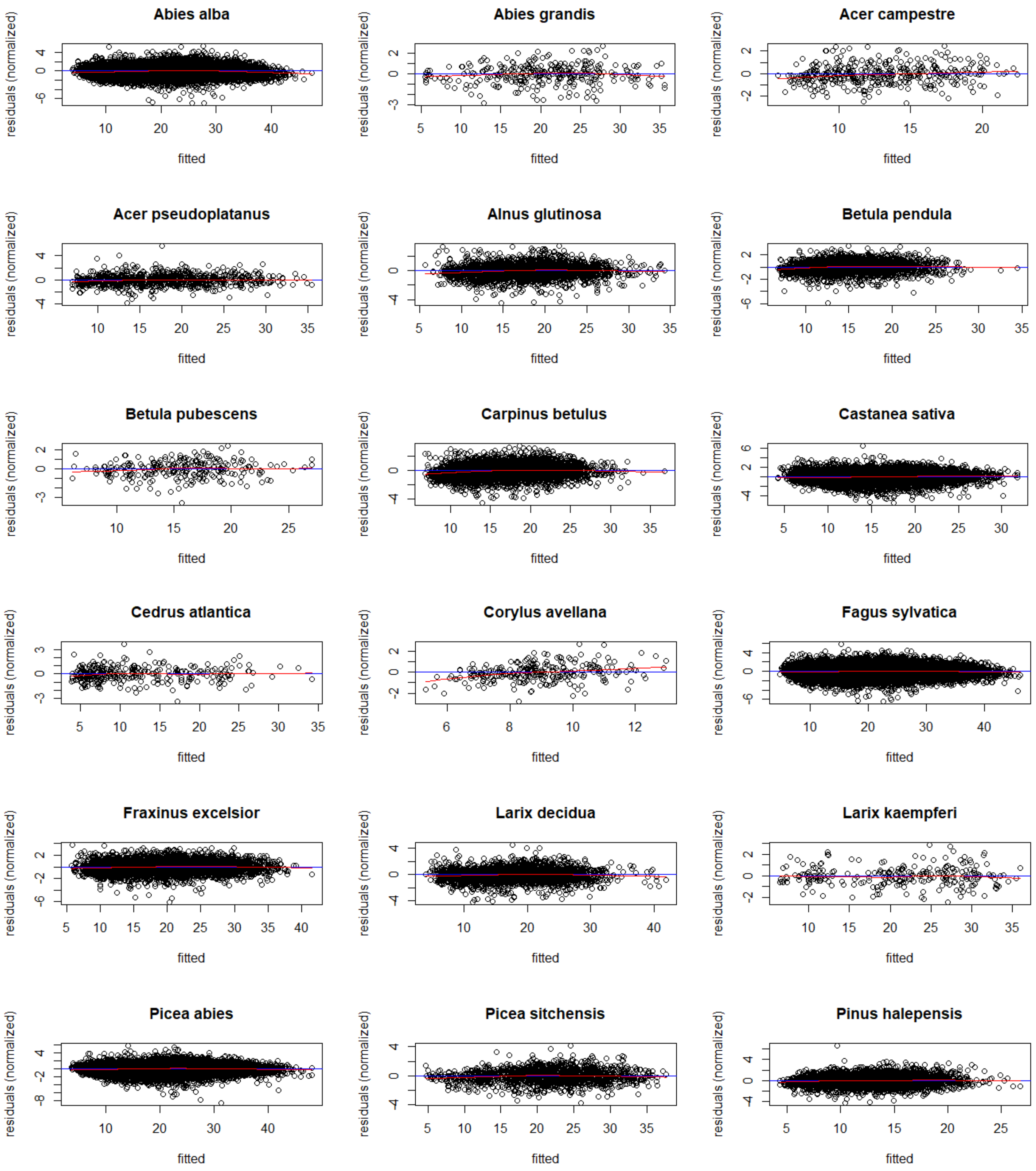

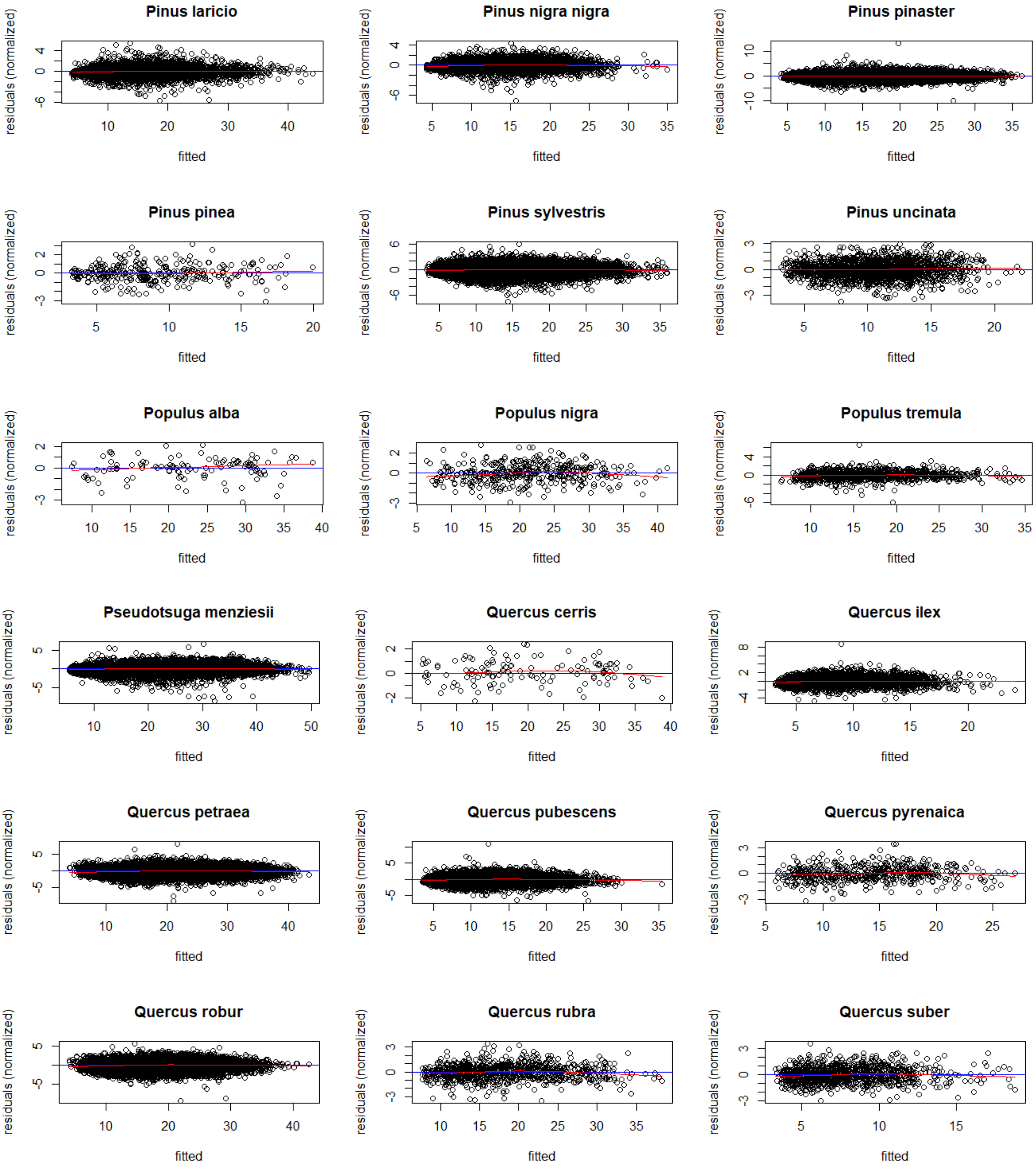
Pinus laricio
Pinus nigra nigra
Pinus pinaster
Pinus pinea
Pinus sylvestris
Pinus uncinata
Populus alba
Populus nigra
Populus tremula
Pseudotsuga menziesii
Quercus cerris
Quercus ilex
Quercus petraea
Quercus pubescens
Quercus pyrenaica
Quercus robur
Quercus rubra
Quercus suber
residuals (normalized)
fitted

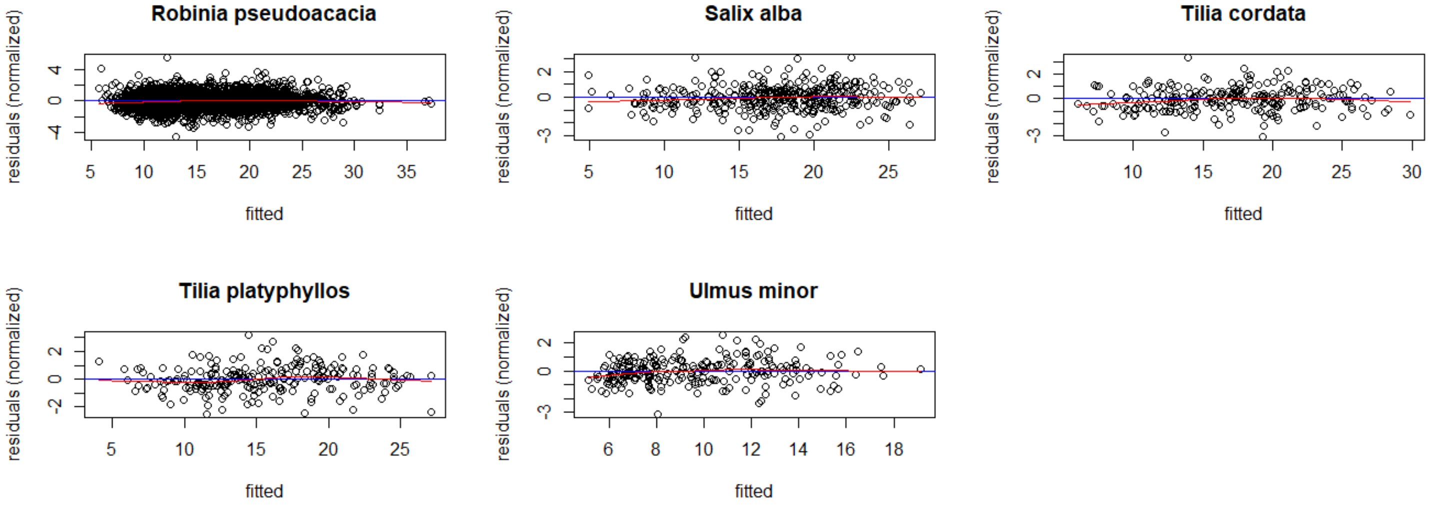
Robinia pseudoacacia
Salix alba
Tilia cordata
Tilia platyphyllos
Ulmus minor
residuals (normalized)
fitted
